# The Human LL-37(17-29) Antimicrobial Peptide Reveals a Functional Supramolecular Nanostructure

**DOI:** 10.1101/2020.02.04.933432

**Authors:** Yizhaq Engelberg, Meytal Landau

**Affiliations:** Department of Biology, Technion-Israel Institute of Technology, Haifa 3200003, Israel; Centre for Structural Systems Biology (CSSB), and European Molecular Biology Laboratory (EMBL), Hamburg 22607, Germany

## Abstract

Protein fibrils that perform biological activities present attractive biomaterials. Here we demonstrate, by crystal structures, the self-assembly of the antibacterial human LL-37 active core (residues 17-29) into a stable structure of densely packed helices. The surface of the fibril encompasses alternating hydrophobic and positively charged zigzagged belts, which likely underlie interactions with and subsequent disruption of negatively charged lipid bilayers, such as bacterial membranes. LL-37_17-29_ correspondingly formed wide, ribbon-like, thermostable fibrils in solution, which co-localized with bacterial cells, and structure-guided mutagenesis analyses supported the role of self-assembly in antibacterial activity. LL-37_17-29_ resembled, in sequence and in the ability to form amphipathic helical fibrils, the bacterial cytotoxic PSMα3 peptide that assembles into cross-α amyloid fibrils. This suggests helical, self-assembling, basic building blocks across kingdoms of life and point to potential structural mimicry mechanisms. The findings offer a scaffold for functional and durable nanostructures for a wide range of medical and technological applications.

## Main

The assembly of basic biological molecules into filamentous structures provides ample opportunities to design bioinspired materials for medical and technological applications^1–8^. One such application is addressing the urgent need to fight microbial aggressive, resistance, infections using materials which allow oral bioavailability, stability in harsh conditions, and long shelf-life. Antimicrobial peptides (AMPs) are canonical components of the innate immune system of many organisms^9^. AMP self-assembly bears functional relevance and can enhance antimicrobial activity^10^. Certain AMPs assemble into well-ordered fibrils that resemble amyloids^11–14^, which are proteins known to form cross-β fibrils composed of tightly mated β-sheets, and have been associated with neurodegenerative and systemic diseases^15,16^. Correspondingly, recent evidence of antimicrobial properties among some human amyloids suggests a potential physiological role of proteins otherwise known as pathological^17–21^.

Our previous findings demonstrated cross-α amyloid fibrillation of the cytotoxic phenol-soluble modulin α3 (PSMα3) peptide secreted by the pathogenic bacterium *Staphylococcus aureus*^22,23^. These fibrils are composed entirely of α-helices that stack perpendicular to the fibril axis into mated ‘sheets’, just as the β-strands assemble in amyloid cross-β fibrils^24^. PSMα3 is toxic to human cells, and some of its truncations and mutants show antibacterial activity^25–27^. Overall, PSMα3 provides a link between toxic activities against human and bacterial cells and unique helical amyloid fibrils^22,23^. Moreover, this architecture offers a scaffold for the design of various supramolecular nanostructures^1,8^. Although originating from different organisms, PSMα3 display sequence similarity with human LL-37 (hLL-37) (Fig. S1), a hCAP-18 protein cleavage product which plays an important role in the first line of defense against pathogens^28^, which is also known to self-assemble^29,30^. Fibrillation of LL-37 was found critical for binding DNA and affecting receptors in the immune system^31^. Both PSMα3 and hLL-37 are cleaved in-vivo into active truncations with diverse activities^32–36^. Some hLL-37 fragments show a diverse array of selectivity against microbial strains, and additional functions within the immune system^34,37–39^. Bacterial proteases can too cleave LL-37, supposedly for degradation or to release virulence factors^36^.

The hLL-37_17-29_ fragment was suggested to serve as the active core of the AMP, showing a different spectrum of antibacterial activity as compared to the full-length protein and other fragments, including being the shortest LL-37 fragment retaining anti-viral activity^36,40–42^. Although not directly detected in-vivo, hLL-37_17-29_ can be cleaved from hCAP-18 or LL-37 by either proteinase K or staphylococcal peptidase I on its N-terminal side, and by trypsin on its C-terminal side. hLL-37_17-29_ is also the region within hLL-37 showing the highest sequence similarity to PSMα3 (Fig. S1), and like the latter, forms a helical monomeric structure shown by NMR experiments^43^. More specifically, hLL-37_17-29_ generates an amphipathic helix with a larger hydrophobic moment compared to the entire hLL-37 and to PSMα3 (Table S1). hLL-37_17-29_ elicited dose-dependent inhibition of Gram-positive *Micrococcus luteus* growth (Fig. S2), with a minimal inhibitory concentration (MIC)^44^ of 22 µM (Fig. 1). It was also active against the *Staphylococcus hominis* bacterium, with a MIC of 39 µM (Fig. S3). We found that hLL-37_17-29_ formed long (several micrometers and longer), ribbon-like, fibrils, visualized using transmission electron microscopy (TEM) (Fig. 2A). Cryogenic electron microscopy (CryoEM) showed that the wide (few hundred nanometers) fibrils are composed of lateral association of thinner fibrils (Fig. S4). The wide fibrils also formed in the presence, and interacted with *M. luteus* cells (Fig. 2B).

**Figure 1.**
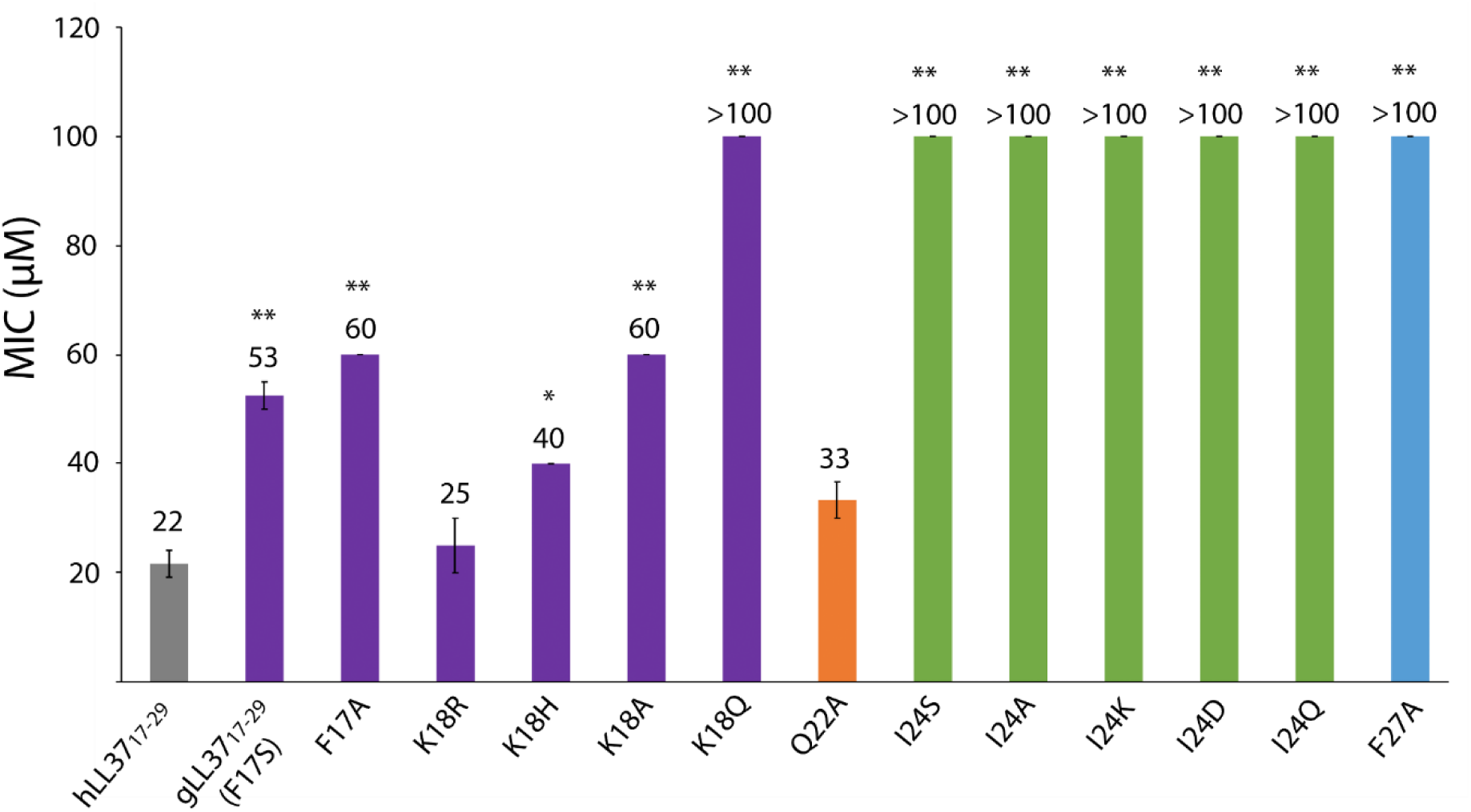
The effect of LL-3717-29 and its mutants on *M. luteus*. hLL-37_17-29_ and some single-point mutants were incubated with *M. luteus* for 24h at a range of concentrations up to 100 µM, and bacterial growth rate was measured by optical density. From the resulting growth curves, the area under the curve (AUC) was calculated. MIC values were defined as the minimal concentration of the peptide which yielded less than 20% of the AUC of the control (bacteria with no added peptides). Mean MIC values are presented above the bars. The MIC of hLL-37_17-29_ is shown by a grey bar. Mutations in residues lining the central pore (Phe17 and Lys18), are shown by purple bars. A mutation in a residue showing minimal contacts with other residues within the fibrillar assembly (Gln22), is shown by an orange bar. Mutations in a residue fully buried in the four-helix bundle (Ile24), are shown by green bars. A mutation in a buried residue (Phe27), contacting other residues within the four-helix bundle and on surrounding helices in the fibrillar assembly, is shown by a blue bar. The experiments were performed at least three times, each on a different day. Error bars indicate the standard error calculated between all measures. A t-test for two-sample assuming equal variances was performed, * indicates p< 5×10^−03^ and ** indicates p< 5×10^−05^, compared to hLL-37_17-29_.

**Figure 2.**
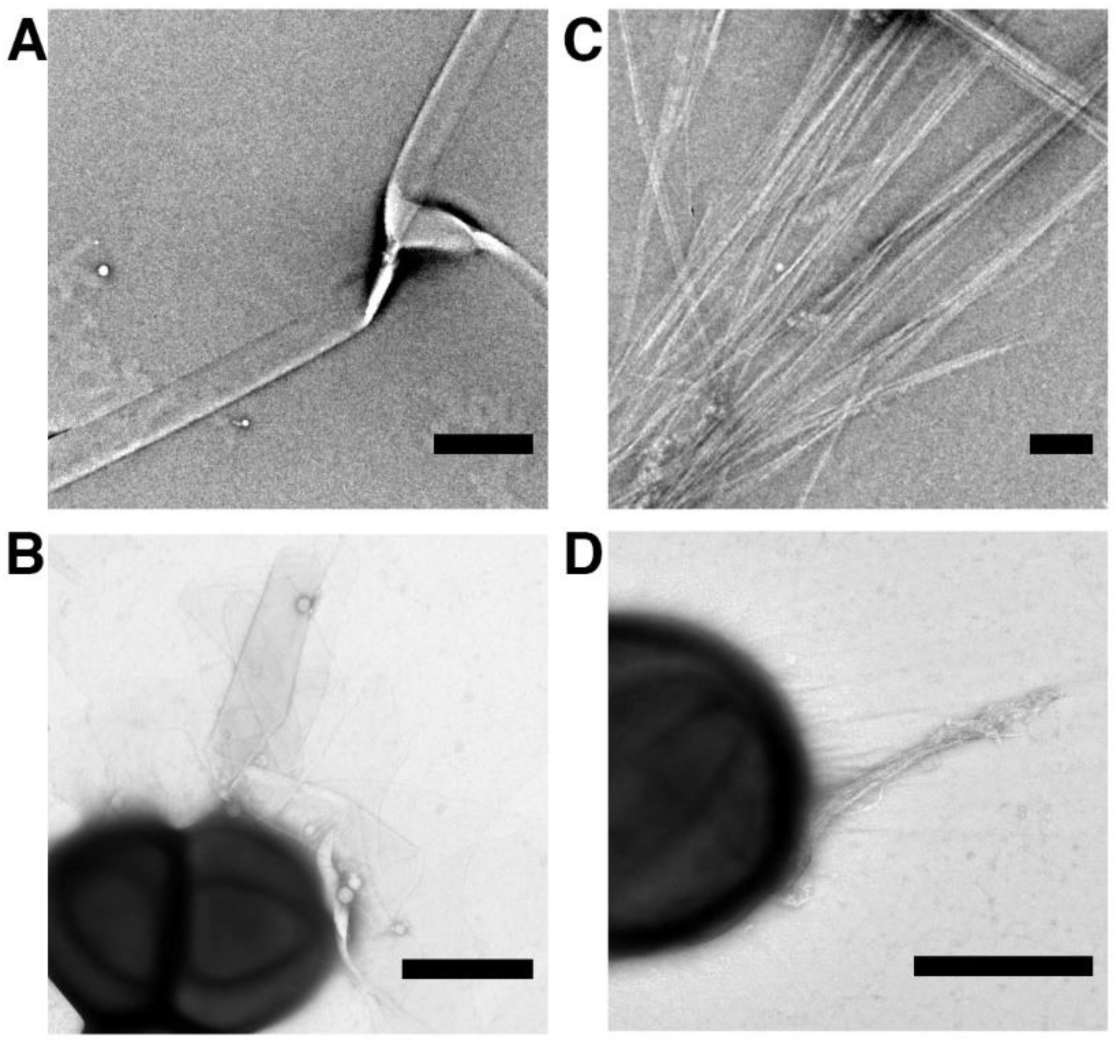
Human and gorilla LL-37_17-29_ fibrillar assemblies and interactions with bacteria. (**A**) An electron micrograph of hLL-37_17-29_ (1 mM) incubated for three days. (**B**) An electron micrograph of hLL-37_17-29_ (30 µM; close to the MIC concentration of 22 µM) incubated with *M. luteus* for 4 h. (**C**) An electron micrograph of gLL-37_17-29_ (1 mM) incubated for three days. (**D**) An electron micrograph of gLL-37_17-29_ (60 µM; around the MIC concentration of 53 µM), incubated with *M. luteus* for 4 h. All scale bars represent 500 nm.

Our determination of the crystal structure of hLL-37_17-29_ at 1.35 Å resolution, revealed self-assembly of amphipathic helices into a densely packed and elongated hexameric structure forming a central pore (Table S2, Fig. 3 and Movie S1). There were two helices in the asymmetric unit of the crystal, with 67% of their individual solvent accessible surface areas buried within the assembly, indicating compact packing. For comparison, in the PSMα3 cross-α structure, each helix is 62% buried in the fibril^22^. In contrast, structures of full-length LL-37, co-crystallized alone or with different lipids, resulted in different levels of assembly, including monomeric, dimeric, tetramers and fiber-like structure of oligomers^45^. The latter was observed when full-length LL-37 was co-crystallized with dodecylphosphocholine (PDB ID 5NNT), showing a repetitive architecture of juxtaposed ‘head-to-tail’ amphipathic helices, with interactions mediated by detergent molecules^45^. This structure formed a much looser packing compared to the LL-37_17-29_ structure, with only 45% of the helix buried within the protein assembly. Overall, the generation of the hLL-37_17-29_ active core allows antibacterial activity along with the formation of highly stable fibrils.

**Figure 3.**
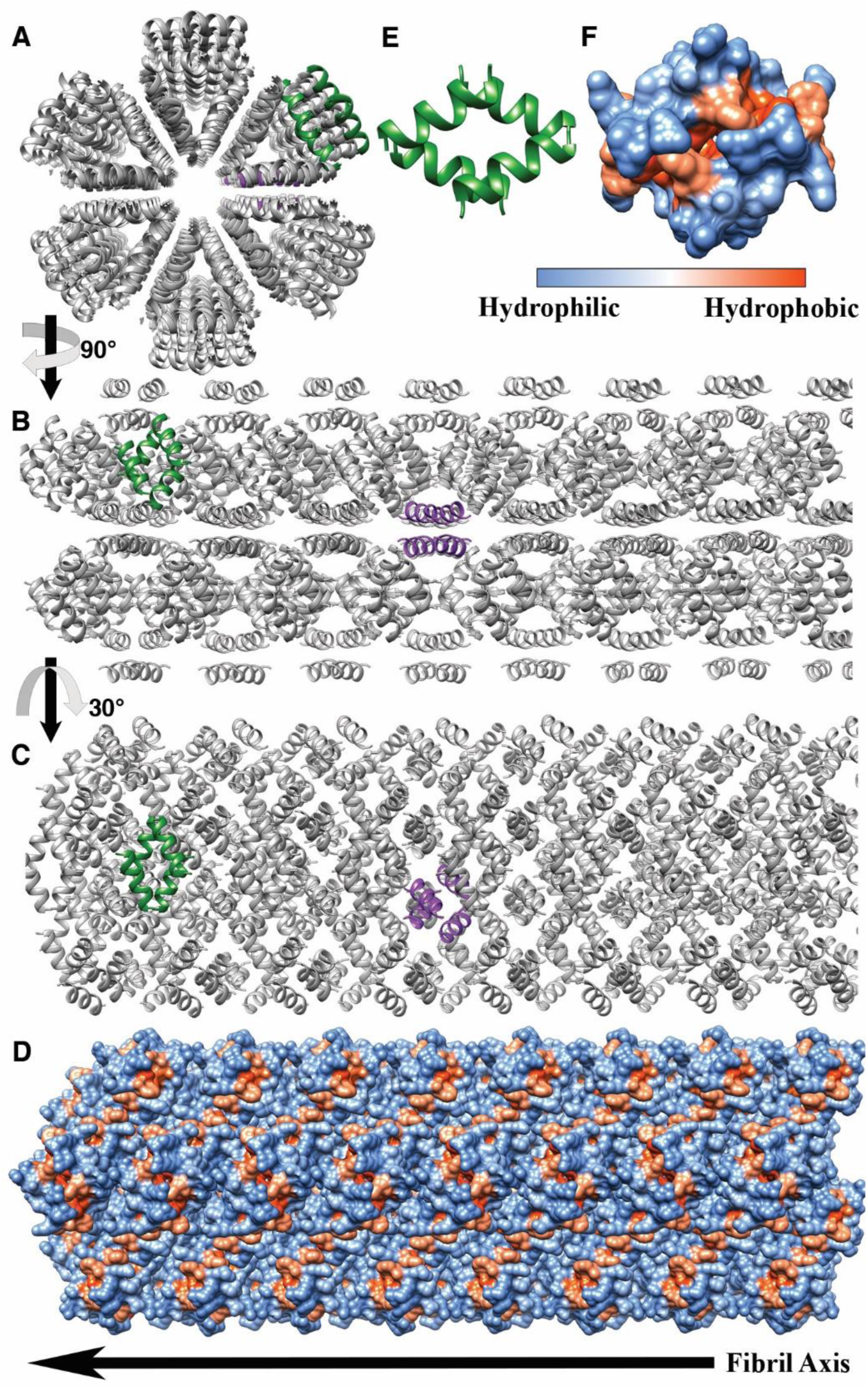
The crystal structure of hLL-37_17-29_

The structure of LL-37_17-29_ lacked amyloid continuous sheets with individual molecules stacked perpendicular to the fibril axis, and correspondingly did not bind the amyloid indicator dye Thioflavin T, in contrast to the cross-α amyloid fibrils of PSMα3^22^ (Fig. S5). Rather, the fibrillar assembly of LL-37_17-29_ was comprised of associating four-helix bundles, each stabilized by a closely packed hydrophobic core (Fig. 3E&F). Arginine residues are lined on the surface of the bundles, with side chains extending outwards, providing an overall highly positively charged molecular electrical potential surface (Fig. S6). The interfaces between bundles comprises a network of polar interactions, including potential salt bridges between Asp26 and Arg23/Arg29 from two adjacent helices, and between Lys25 and the C-terminus of an adjacent helix (Fig. S7). Due to the symmetry in the structure, each bundle of four helices could form sixteen inter-helical polar interactions with adjacent helices in the assembly (Fig. S7). In addition, Asp26 could potentially form a salt bridge with Arg29 on the same helix. Such intra-helical salt-bridges are associated with increased α-helical stability^46^. Phe11, facing towards another Phe11 residue from an adjacent helix, contributes to hydrophobic packing between bundles.

The overall stable assembly of LL-37_17-29,_ which includes a network of polar interactions and hydrophobic packing, corresponded with the thermostability of the fibrils, as visualized by electron micrographs after heating to 60°C or 80°C (Fig. S8). Some disassembly at the edge of the wide fibrils, into thinner fibrils, could be observed after the 80°C heat shock, or after further incubation following the 60°C heat shock exposing a lateral fibril association. In comparison, collagen fibrils, which provides physical support to tissues, disintegrate upon heating to 65°C^47^. In addition, the surface charge and colloidal stability of the LL-37_17-29_ were assessed using zeta potential measurements^48^, showing concentration dependent values, reaching +25 mV at 1 mM (Table S3). Congruently, other positively charged biological polymers were previously proposed for antimicrobials and drug delivery applications^49,50^.

The fibrillar assembly in the crystal structure created alternating hydrophobic and polar (positively charged) zigzagged belts on its surface (Fig. 3D), suggesting interactions with and disruption of negatively charged lipid bilayers, such as bacterial membranes^51^. Confocal microscopy images of fluorescein isothiocyanate (FITC)-labeled hLL-37_17-29_, which showed antibacterial activity similar to that of the unlabeled peptide (Fig. S9), confirmed aggregation and co-localization of hLL-37_17-29_ with bacterial cells (Fig. S10 and Movie S2). The role of the self-assembly in the antibacterial activity of hLL-37_17-29_ was examined using single-point mutations designed based on the solved crystal structure. Alanine substitution of Ile24, the most buried residue within the core of the four-helix bundle (Fig. S11 and Table S4), abolished antibacterial activity against *M. luteus* (Fig. 1) and against *S. hominis* (Fig. S3). This mutation failed to exhibit peptide assembly around bacterial cells (Fig. S12). Correspondingly, confocal microscopy images of the FITC-labeled I24A mutant, which, like the unlabeled peptide, was also ineffective against *M. luteus* (Fig. S9), indicated absence of peptide aggregation (Fig. S10). Likewise, the I24S, I24K, I24Q and I24D mutations all abolished antibacterial activity against *M. luteus* (Fig. 1), and confocal microscopy images of the FITC-labeled I24S inactive mutant (Fig. S9) showed no detectable aggregation (Fig. S10). Zeta potential measurements of the I24A inactive mutant showed significantly lower values compared to the native sequences, reaching only +14 mV at 1 mM (Table S3). Since I24A is a conservative substitution with no change in net charge, we attribute the lower zeta potential values to lack of an organized particle assembly by the mutant.

Phe27 is another residue significantly buried within the fibrillar assembly (Table S4), forming contacts with residues on the four-helix bundle and with other helices in the fibrillar assembly (Fig. S11). The F27A mutation abolished antibacterial activity against *M. luteus* (Fig. 1) and failed to aggregate when incubated with the bacteria (Fig. S12). In contrast to the buried Ile24 and Phe27, Gln22 was the least buried residue in the assembly, forming minimal contacts within adjacent helices (Fig. S11 and Table S4). Consequently, the Q22A mutation resulted in minimal change in activity against *M. luteus* (a MIC of 33 µM; Fig. 1) and was also active against *S. hominis*, with a MIC of 53 µM (Fig. S3). Electron micrographs of the Q22A mutant showed large fibrous nanostructures contacting the bacterial cells (Fig. S12). Correspondingly, confocal microscopy images of the FITC-labeled Q22A, which was slightly less active than the unlabeled Q22A peptide (Fig. S9), formed aggregates which co-localized with the bacterial cells (Fig. S10). Overall, the mutagenesis analyses indicated the importance of hLL-37_17-29_ self-assembly in its antibacterial activity and in direct interactions with bacterial cells, and supported a plausible active fibril arrangement.

Investigation of the N-terminal residues which are not buried within the assembly (Table S4) but which face the central pore, showed that the F17A and K18A mutants display a similar reduction in antibacterial activity against *M. luteus*, with a MIC of 60 µM (Fig. 1). Maintaining the positive charge via a K18R substitution, showed a very similar MIC to that of hLL-37_17-29_, while the K18H substitution showed slightly reduced activity (Fig. 1). In contrast, substitution to the polar but uncharged glutamine (K18Q) fully abolished activity (Fig. 1). The results suggest that the two residues facing the pore are important for activity, with the positive charge of Lys18 being the critical determinant. We therefore cannot conclude about the specific role of the central pore in activity, as the effect of substitutions might be related to the reduced positive charge^52,53^ regardless of its structural location.

The gorilla LL-37 (gLL-37) sequence contains two amino acid substitutions when compared to hLL-37, with one at position 17, the first residue of LL-37_17-29_, substituting phenylalanine with serine (corresponding to a F17S mutation). gLL-37_17-29_ exhibited slightly weaker antibacterial activity against *M. luteus* compared to hLL-37_17-29_, with a MIC of 53 µM (Fig. 1), similar to the effect of the F17A mutant. The 1.1 Å resolution crystal structure of gLL-37_17-29_ displayed a similar assembly compared to hLL-37_17-29_ (Table S2 and Fig. S13), with almost identical backbone positions and root-mean-square deviation (RMSD) of 0.15 Å, differing only in the two N-terminal positions that lined the central pore (Fig. S14). Accordingly, both hLL-37_17-29_ and gLL-37_17-29_ formed wide fibrillary structures, as observed by cryogenic electron micrographs (Fig. S4), and engaged in direct contact with *M. luteus* cells (Fig. 2).

To conclude, the atomic structures of human and primate antibacterial LL-37_17-29_ showed a functional supramolecular nanostructure of densely packed amphipathic helices. This assembly into stable fibrils with a surface forming hydrophobic/charged zigzagged belts can be used as scaffolds for wide-ranging applications in bio and nanotechnology, regenerative medicine and bioengineering^54^, with the invaluable advantage of an inherent antibacterial activity. Links between fibril formation and antimicrobial activity are accumulating^11–14,17–21,55^, and here, we provide atomic-level insight for such example. Further elucidation of the interplay between antimicrobial activity and fibril formation and morphology will aid the design of AMPs with enhanced potency, selectivity, stability, bioavailability and shelf-life. Successful design of such functional nanostructures with tunable self-assembly might provide novel antibacterial therapeutics or coating of medical devices, and will may target other roles of AMPs in immunomodulation and in killing cancerous cells^56^.

The LL-37_17-29_ structure differs from known helical fibrils such as the toxic cross-α amyloid fibrils of PSMα3, and from structural fibrils such as collagen, actin, and fibrinogen. It overall presents a type of self-assembly which, to the best of our knowledge, is distinct from other protein fibrils, with a role in direct killing of bacterial cells still to be fully determined. Despite the different arrangement, the sequence similarity between the human LL-37 and the bacterial PSMα3, and their shared ability to form helical functional fibrils, suggest a possible molecular or structural mimicry mechanism used by the bacteria to provide immune-evasive and survival strategies^57,58^. This also points to potential functional building blocks across kingdoms of life in the form of densely packed amphipathic helical fibrils, complementing the exciting hypotheses about short amyloid peptides serving as prebiotic information-coding molecules^59–61^.

The crystal structure of hLL-37_17-29_, determined at 1.35 Å resolution. The crystal packing shows self-assembly of amphipathic helices into a densely packed, elongated hexameric fibril with a central pore. The fibril is composed of four-helix bundles with a hydrophobic core that associated via a network of polar interaction (Fig. S7). (**A**) The assembly is shown as grey ribbons, with two representative four-helix bundles colored green and purple to emphasize orientation in the fibril. (**B**) The view is rotated by 90° in relation to panel A, showing the structure along the fibril axis. (**C**) The view is rotated by 30° along the fibril axis compared to panel B. (**D**) The fibril, in the same orientation as in panel C, is shown in a surface representation, colored by hydrophobicity, according to the scale bar. (**E**) An isolated four-helix bundle shown as green ribbons. (**F**) The four-helix bundle, in the same orientation as in panel E, shown in a surface representation colored by hydrophobicity, according to the scale bar.

## Acknowledgments

We are grateful for Duilio Cascio (University of California Los Angeles) for advice and help with crystal structure determination. We thank Einat Netzer for help in conducting experiments, Carolin Seuring for advice on microscopy images, and Eilon Barnea, Nir Salinas and Yael Pazy-Benhar for commenting on the manuscript. We acknowledge guidance and technical support provided by Yael Pazy-Benhar and Dikla Hiya at the Technion Center for Structural Biology (TCSB), Na’ama Koifman and Ellina Kesselman from the Russell Berrie Electron Microscopy Center of Soft Matter, Yael Lupo-Haber and Nitsan Dahan from the Life Science and Engineering Infrastructure Center, all at the Technion, Israel. This research was supported by Israel Science Foundation (grant no. 560/16), Israel Ministry of Science, Technology & Space (grant no. 78567), BioStruct-X, funded by FP7, and the iNEXT consortium of Instruct-ERIC. The synchrotron MX data collection experiments were performed at beamlines ID23-EH2 at the European Synchrotron Radiation Facility (ESRF), Grenoble, France, and at beamline P14, operated by EMBL Hamburg at the PETRA III storage ring (DESY, Hamburg, Germany). We are grateful to the teams at ESRF and EMBL Hamburg. We would like to thank Thomas R. Schneider and Gleb Bourenkov for their assistance in operating the P14 beamline.

---

**Figure S1.**
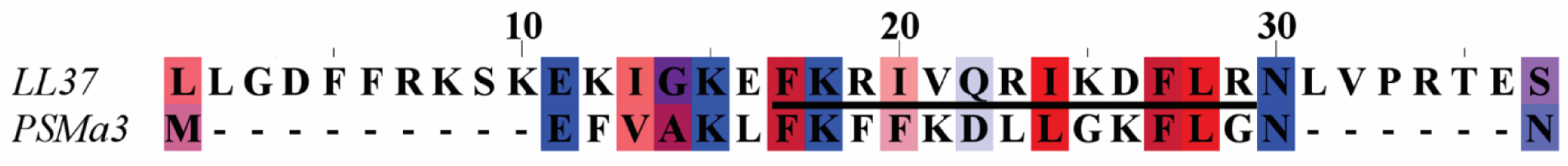
Sequence alignment of human LL-37 and bacterial PSMα3. Sequence alignment between human LL-37 (UniProt ID P49913) and *S. aureus* PSMα3 (UniProt ID H9BRQ7). Amino acids are color-coded by their physicochemical properties^62^. Identity and similarity between the two sequences were 19% and 24%, respectively. The hLL-37_17-29_ segment within the full sequence of hLL-37 is underlined and constitutes the most conserved region between the two peptides, with 31% identity and 39% similarity to the equivalent segment in PSMα3.

**Figure S2.**
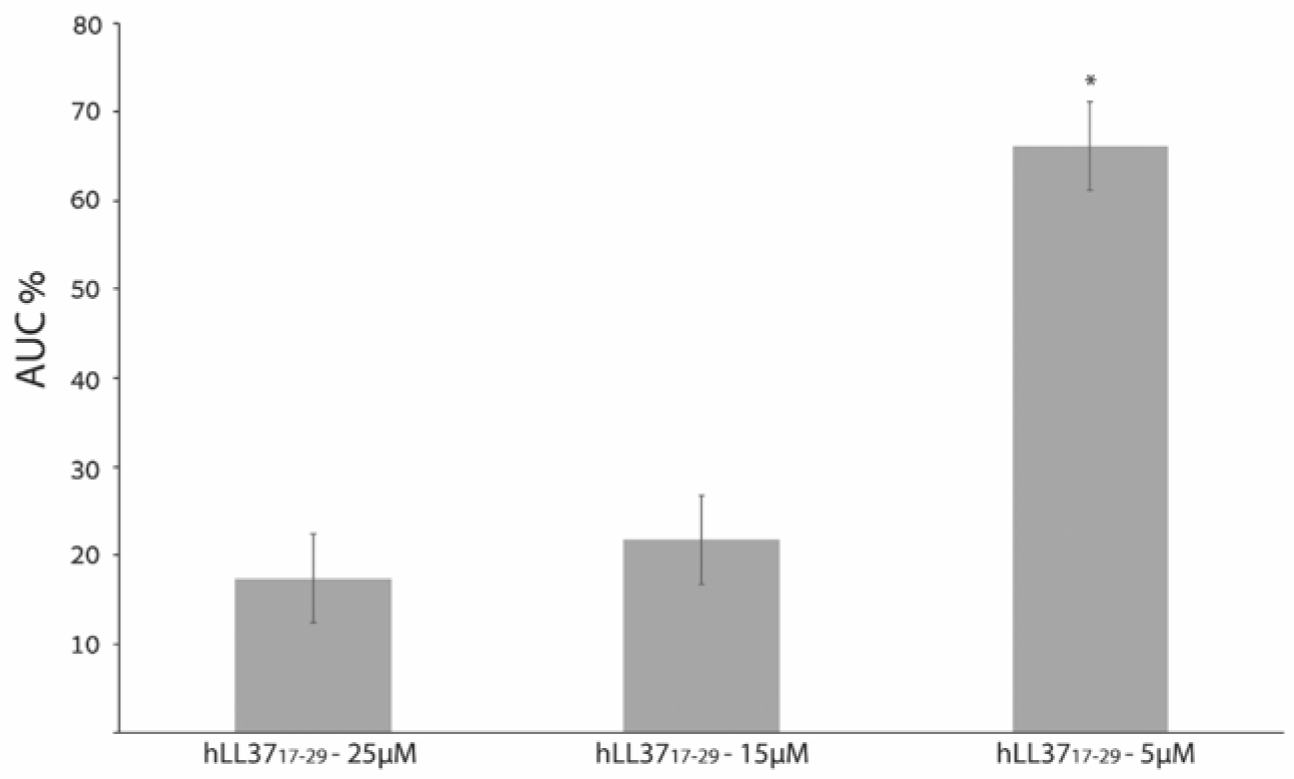
Human LL-3717-29 concentration-dependent inhibition of *M. luteus* growth. The integrated area under the curve (AUC) of bacterial growth over 24h is presented as percentage of control (peptide-free samples). All experiments were performed in triplicates, which were averaged. The experiments were performed at least three times, on different days. Error bars represent the standard deviation of the mean (from the averaged triplicates of all biological repeats) and divided by the root of the number of repeats. For significance, a paired two-sample t-test, assuming equal variances, was performed; *indicates p<5×10^−05^ compared to hLL-37_17-29_ 25µM.

**Figure S3.**
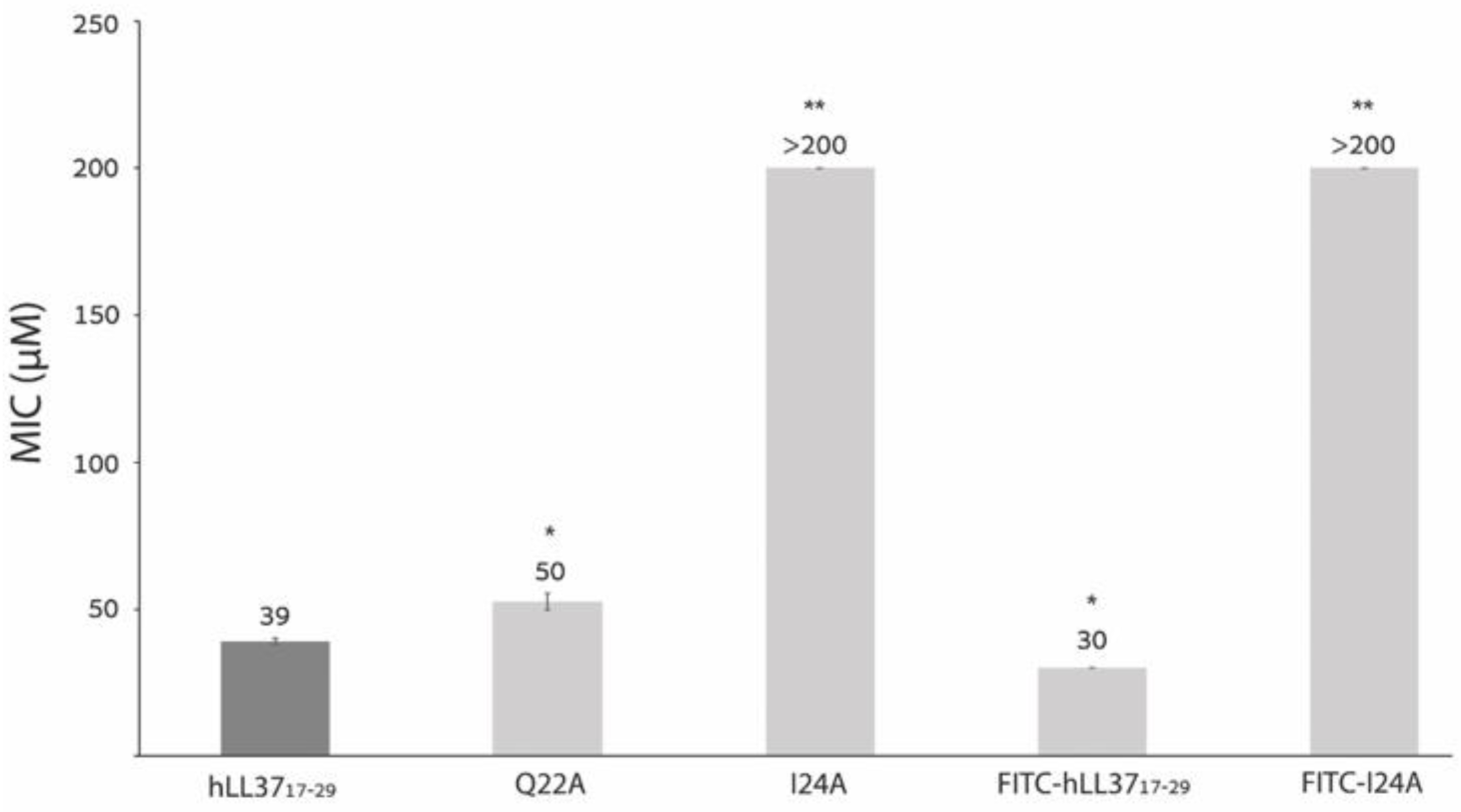
Inhibition of *S. hominis* growth by different hLL-3717-29 mutants and FITC-conjugates. Growth inhibition of *S. hominis* by different hLL-37_17-29_ mutants and FITC-conjugates, as expressed by the MIC values indicated above the bars. The highest tested concentration of the peptides was 200 µM. All experiments were performed in triplicates, which were averaged. The experiments were performed at least three times, on different days. Error bars represent the standard deviation of the mean (from the averaged triplicates of all biological repeats). A paired, two-sample Student’s t-test, assuming equal variances, was performed; * indicates p<0.001, ** indicates p<5×10^−10^ compared to hLL-37_17-29_.

**Figure S4.**
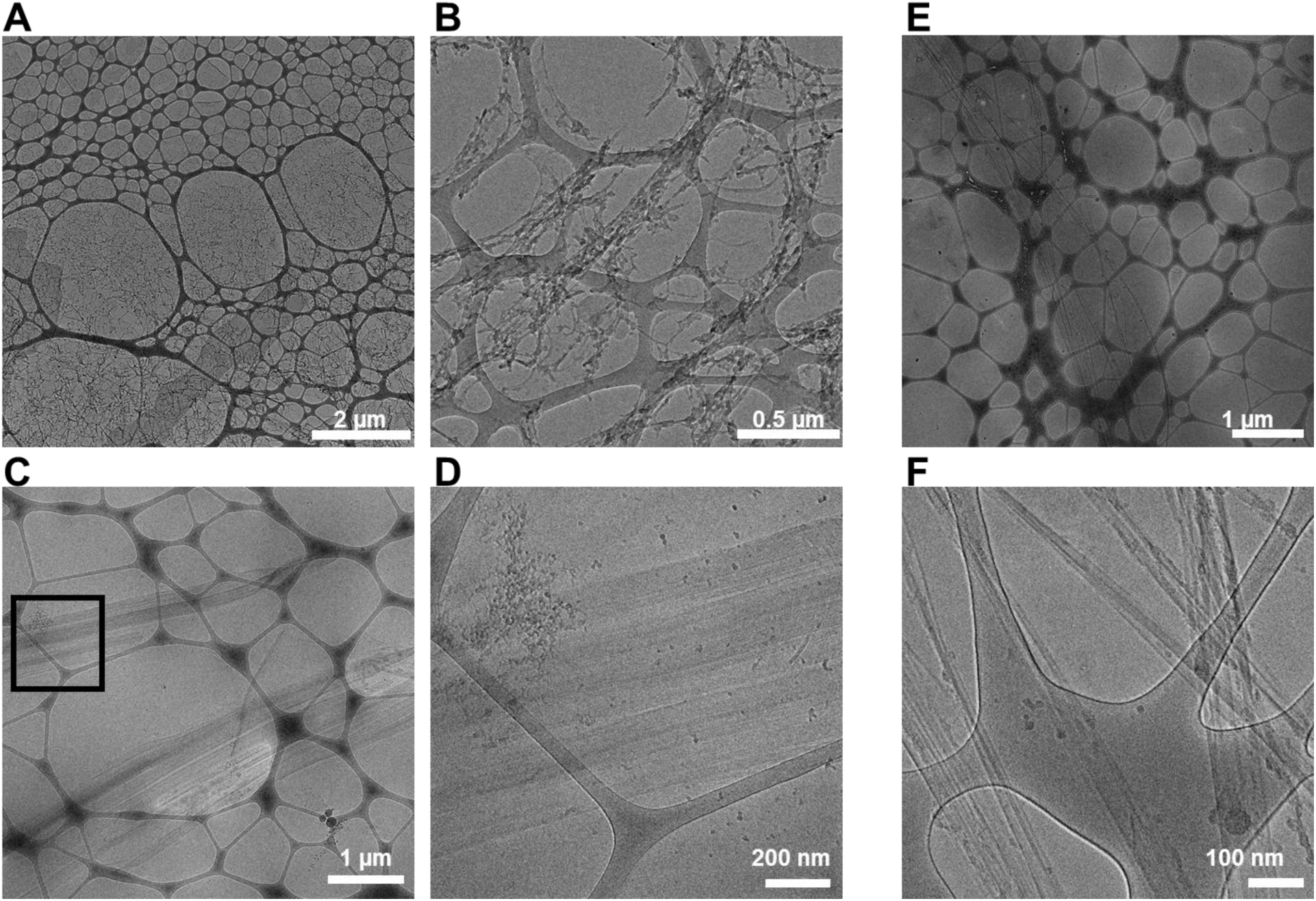
Cryo-electron micrographs of human and gorilla LL-3717-29. Cryogenic electron micrographs of human (**A-D**) and gorilla (**E-F**) LL-37_17-29_. (**A-B**) Micrographs at two different magnifications show massive fibrillation of 5 mM hLL-37_17-29_, after incubation at 37 °C, for three days. (**C-D**) hLL-37_17-29_ (2 mM) was incubated with 2.7 mM SDS for 10 days, at 37 °C. The image in panel D is a zoom-in view of the boxed image in panel C. The micrographs display the formation of straight fibrils that bundle to form several hundred nanometer-wide ribbons, similar to what was observed in the negative-staining TEM images (Fig. 2). (**E-F**) Micrographs of 1 mM gLL-37_17-29,_ incubated for three days, at 37 °C, show the formation of very long (several micrometers) and straight fibrils that also bundle into wide ribbon-like fibrils.

**Figure S5.**
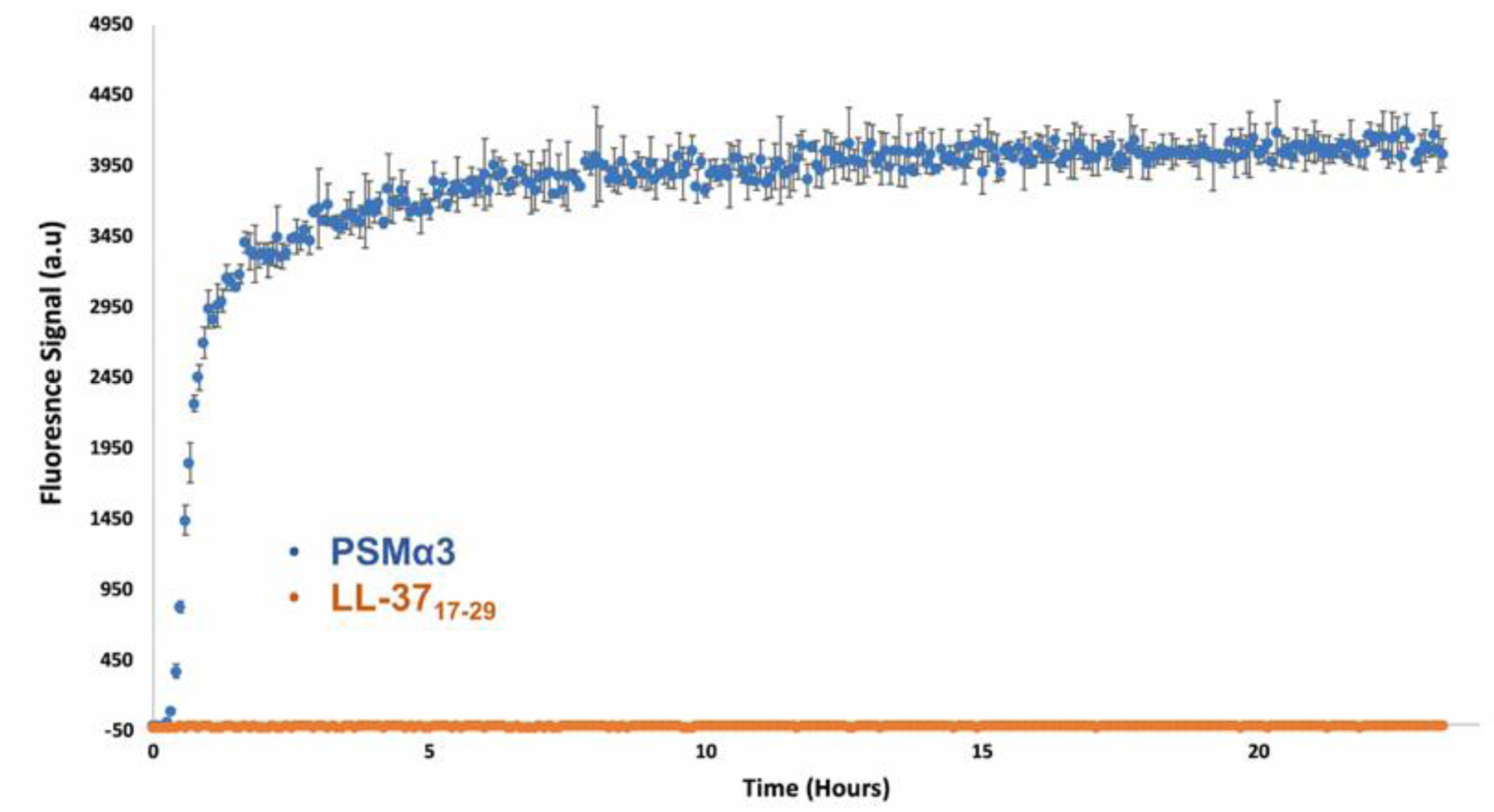
ThT fluorescence kinetics of hLL-3717-29 and PSMα3. ThT fluorescence kinetics in the presence of 1 mM hLL-37_17-29_ (orange curve) or 50 µM PSMα3 (blue curve). PSMα3 showed ThT binding, indicating rapid fibril formation, while hLL-37_17-29_ failed to bind ThT. Measurements were performed in triplicates and values were averaged, appropriate blanks were subtracted, and the resulting values were plotted against time. Error bars represent standard errors of the mean. The entire experiment was repeated at least three times, on different days, showing similar results.

**Figure S6.**
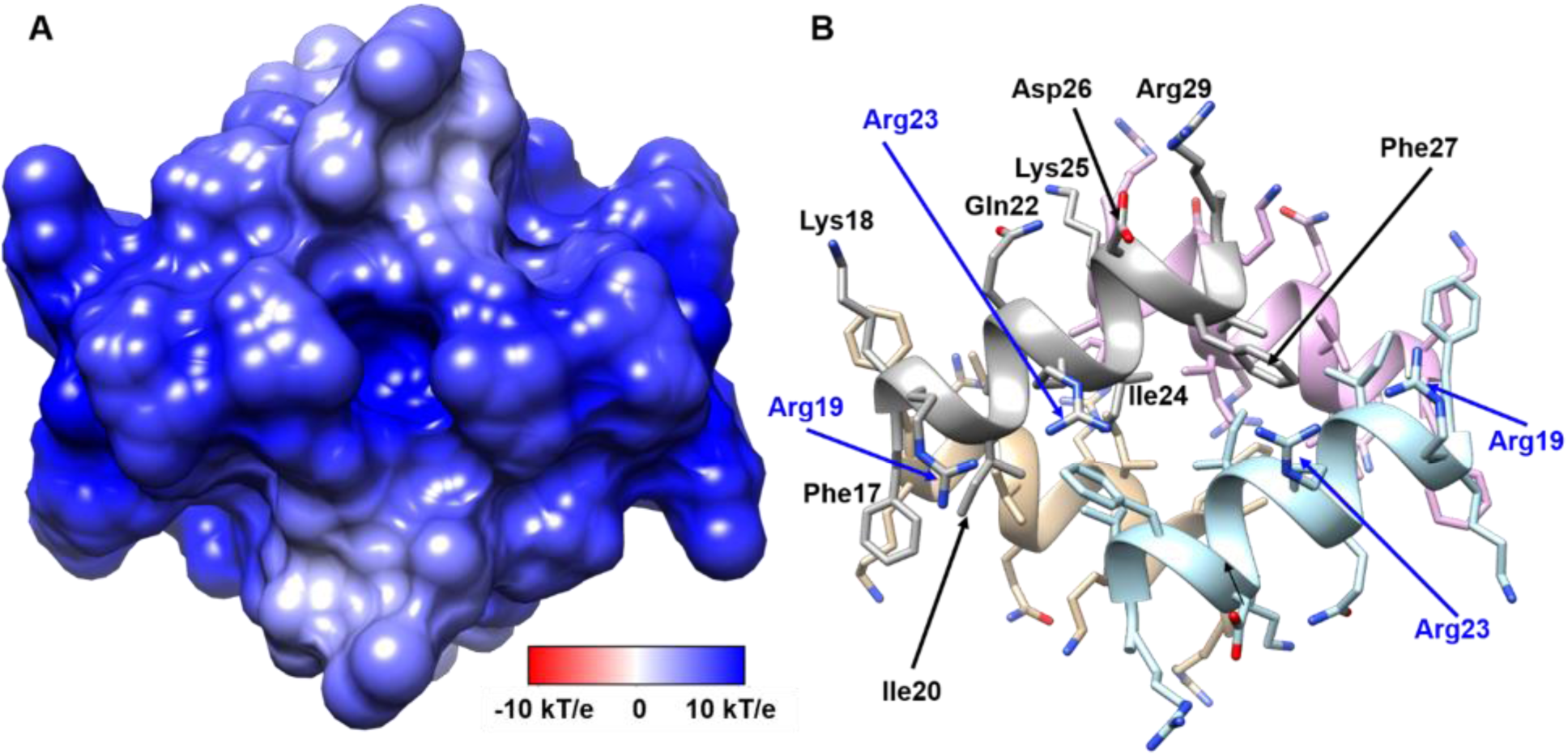
Positively charged electrostatic surface of the four-helix bundle of hLL-37_17-29_. (**A**) A projection of the electrostatic potential (φ) onto the molecular surface of the four-helix bundle of hLL-37_17-29_; the scale bar indicates φ ranges between −10 kT/e (dark red) and 10 kT/e (dark blue). (**B**) The bundle is displayed as ribbons, in the same orientation as in panel A, with side chains shown as sticks. The ribbons and carbons of each of the four helices are colored differently (grey, light blue, tan and pink) and non-carbon atoms are colored by atom type (oxygen in red and nitrogen in blue). Residues are labeled. Arg19 and Arg23 (labeled blue) from two helices lie across each side of the bundle.

**Figure S7.**
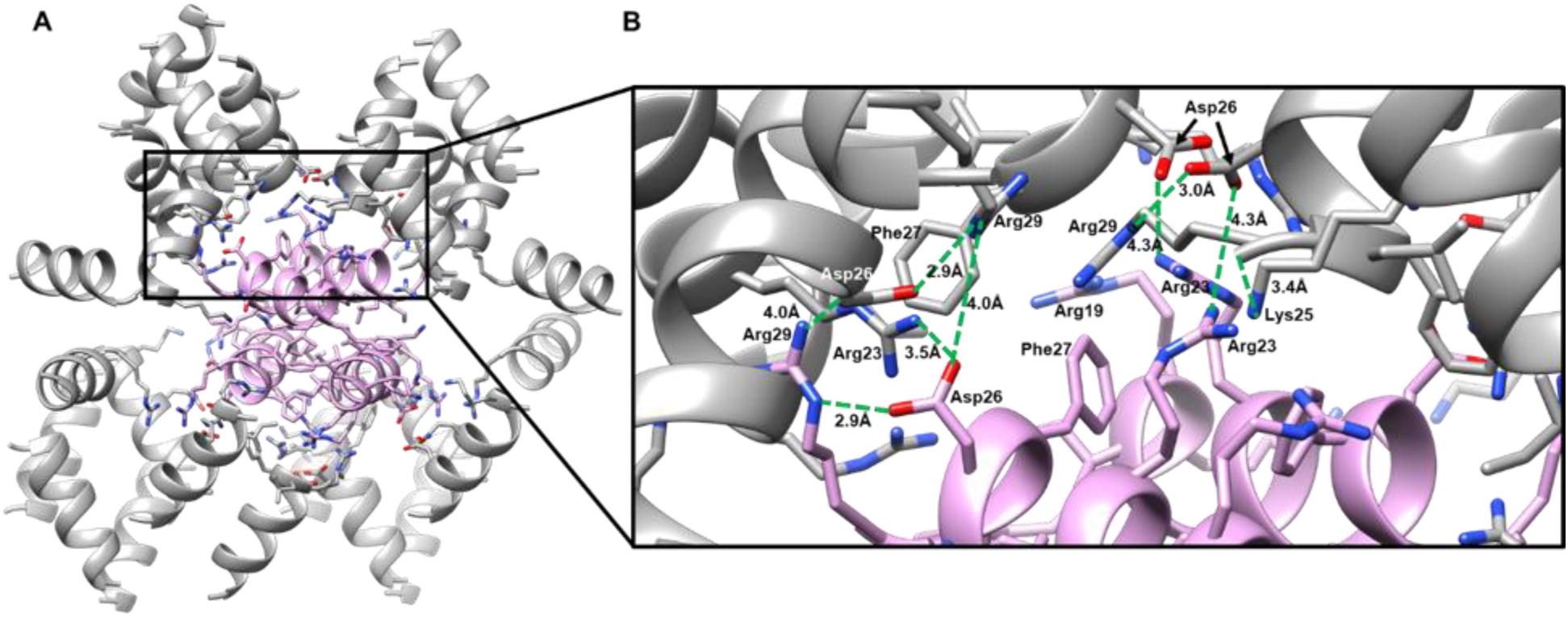
Interfaces of the four-helix bundle with surrounding helices in the fibril assembly. **(A**) One representative four-helix bundle is shown by pink ribbons, and surrounding helices are colored grey. Side chains of the “pink” bundle and residues of surrounding helices which contact the bundle are shown as sticks, colored by atom type (oxygen in red and nitrogen in blue). The asymmetric unit of the crystal contains two chains which are almost identical (RMSD of 0.13 Å), thus, the bundle shows almost four identical interfaces. (**B**) A zoom-in view. Potential polar interactions are displayed, and distances between residues are indicated: Asp26 can form inter-helical electrostatic interactions with Arg23, and with Arg29 from a different helix, and intra-helical electrostatic interactions with Arg29. In addition, Lys25 can form electrostatic interactions with the negatively charged C-terminus of an adjacent helix. Overall, each helix shows four inter-helical and one intra-helical electrostatic interactions. In addition, Phe27 faces the middle of the interface, contributing to hydrophobic packing.

**Figure S8.**
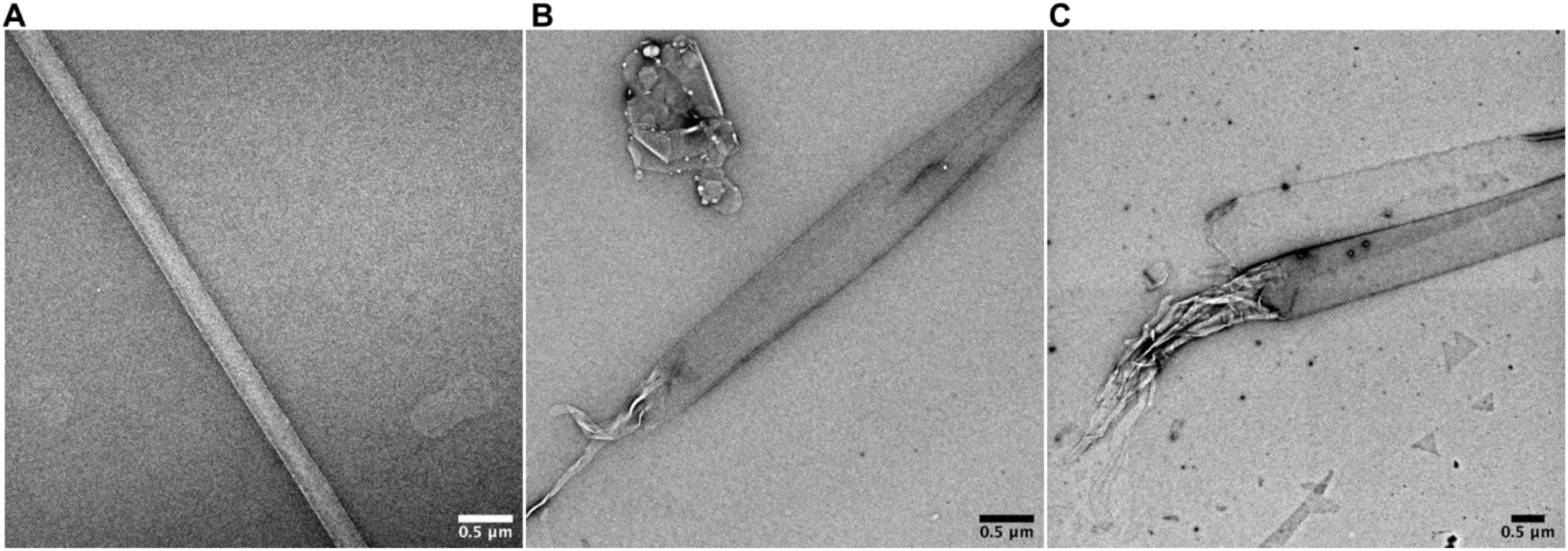
Thermostability of hLL-37_17-29_ fibrils. Transmission electron micrographs of 2mM LL-37_17-29_ incubated for 3 days. (**A**) The sample was heated to 60°C for 10 min. (**B**) The sample was incubated for additional 24hr at 37°C after the 60°C heat shock. (**C**) The sample was heated to 80°C for 10 min.

**Figure S9.**
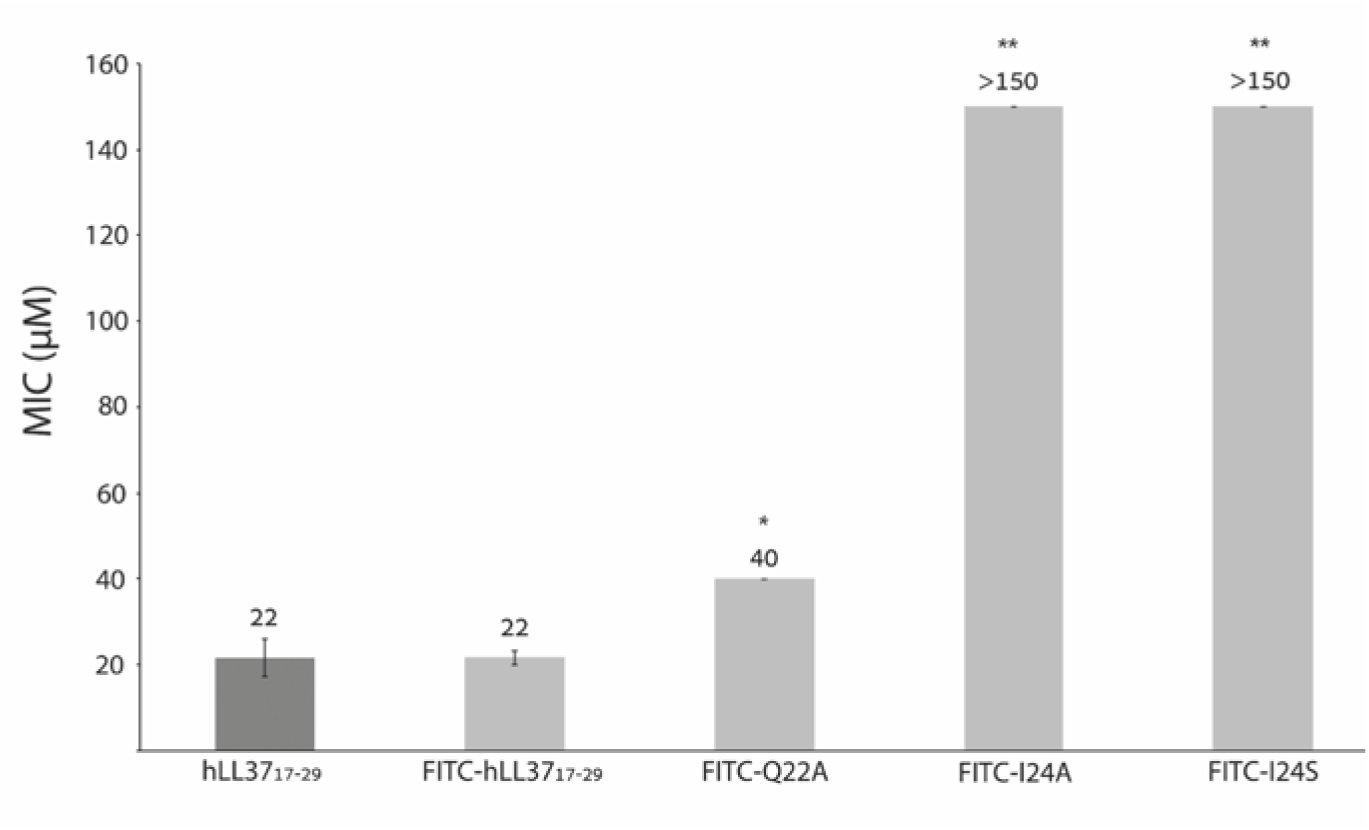
Growth inhibition of *M. luteus* by FITC-labeled LL-3717-29 derivatives. Growth inhibition of *M. luteus* by FITC-hLL-37_17-29_ and its mutants is displayed as mean MIC values provided above the bars calculated as described before. The highest tested concentration of the peptides was 150 µM. All experiments were performed in triplicates, which were averaged. The experiments were performed at least three times, on different days. Error bars represent the standard deviation of the mean (from the averaged triplicates of all biological repeats). A paired, two-sample Student’s t-test assuming equal variances was performed; * indicates p< 5×10^−03^ and ** indicates p< 5×10^−07^ compared to hLL-37_17-29_.

**Figure S10.**
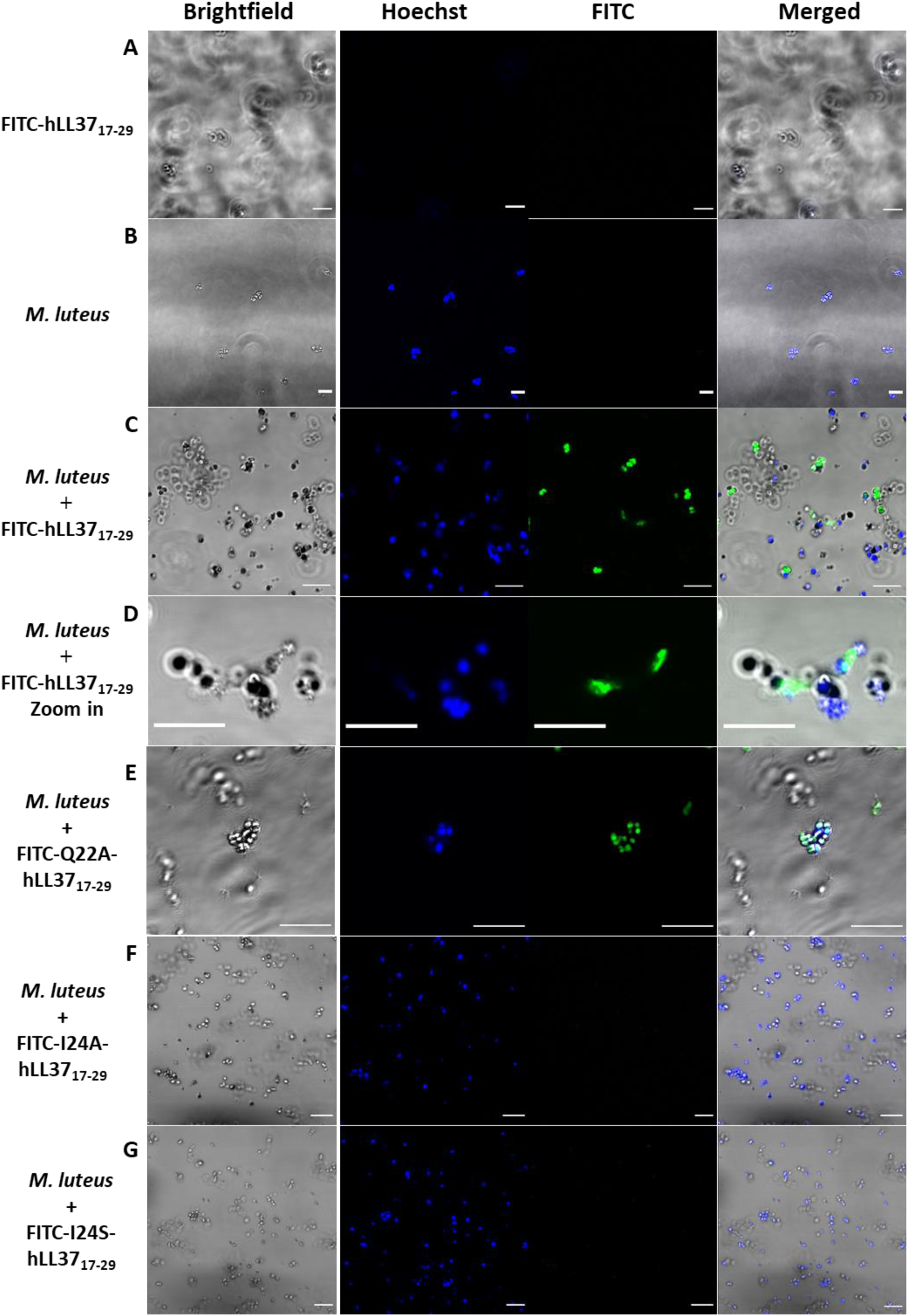
Confocal microscopy images of *M. luteus* incubated with FITC-LL-37_17-29_ and its mutants. Representative confocal microscopy images of *M. luteus* incubated for 4 h with FITC-labeled LL-37_17-29_ or mutants. The FITC channel was merged with the bright-field channel to show peptide location with respect to the bacteria (right column). A 20 µm scale bar is shown for all images. (**A**) A control sample containing 200 µM FITC-hLL-37_17-29_ with no bacterial cells, showing no visible fluorescent signals. (**B**) A control sample containing *M. luteus* with no peptide, showing the blue stained bacterial cells. (**C-F**) *M. luteus* with: (**C-D**) 30 µM FITC-hLL-37_17-29_ and a zoom-in view in panel D, (**E**) 50 µM FITC-hLL-37_17-29_ Q22A, (**F**) 150 µM FITC-hLL-37_17-29_ I24A or (**G**)150 µM FITC-hLL-37_17-29_ I24S. The FITC-hLL-37_17-29_ (**C-D**) and FITC-hLL-37_17-29_ Q22A (**E**) aggregated (bright green foci) and co-localized with the bacterial cells. The FITC-hLL-37_17-29_ I24A (**F**) and I24S (**G**) inactive mutants (Fig. 1) do not undergo aggregation.

**Figure S11.**
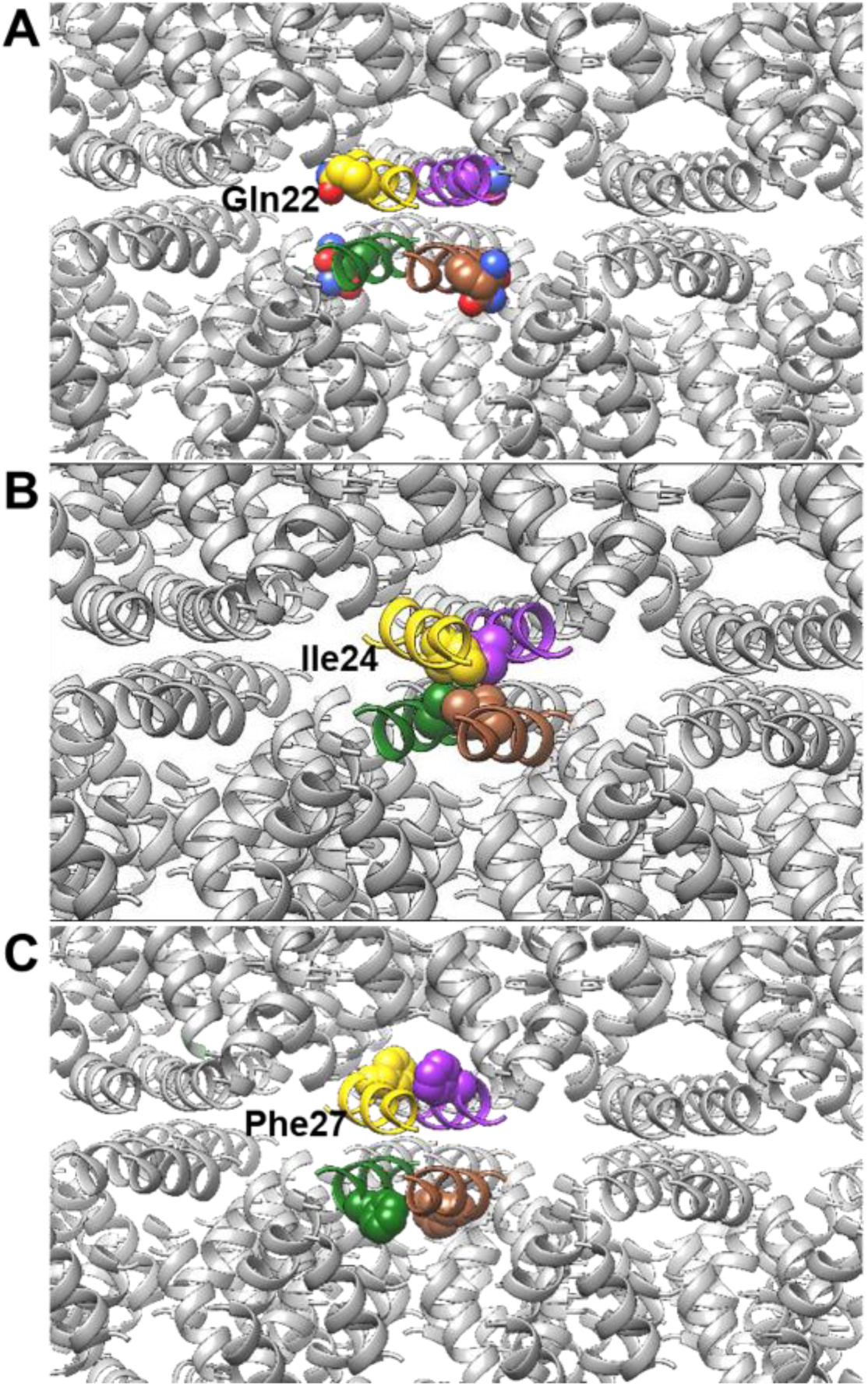
Structural location of mutated LL-37_17-29_ residues. A zoom-in view of the fibrillar assembly, focusing on one four-helix bundle, with each helix colored differently. In each panel, residues that were substituted in our assays are individually shown using space-filling model. (**A**) Gln22 faces outward from the bundle, forming very few contacts with adjacent helices, with only 13% of its SASA buried in the assembly (Table S3). (**B**) Ile24 is completely buried (95% of its SASA) inside the four-helix bundle. (**C**) Phe27 faces away from the bundle yet contacts both other residues on the same bundle and adjacent helices, with 85% of its SASA buried in the assembly.

**Figure S12.**
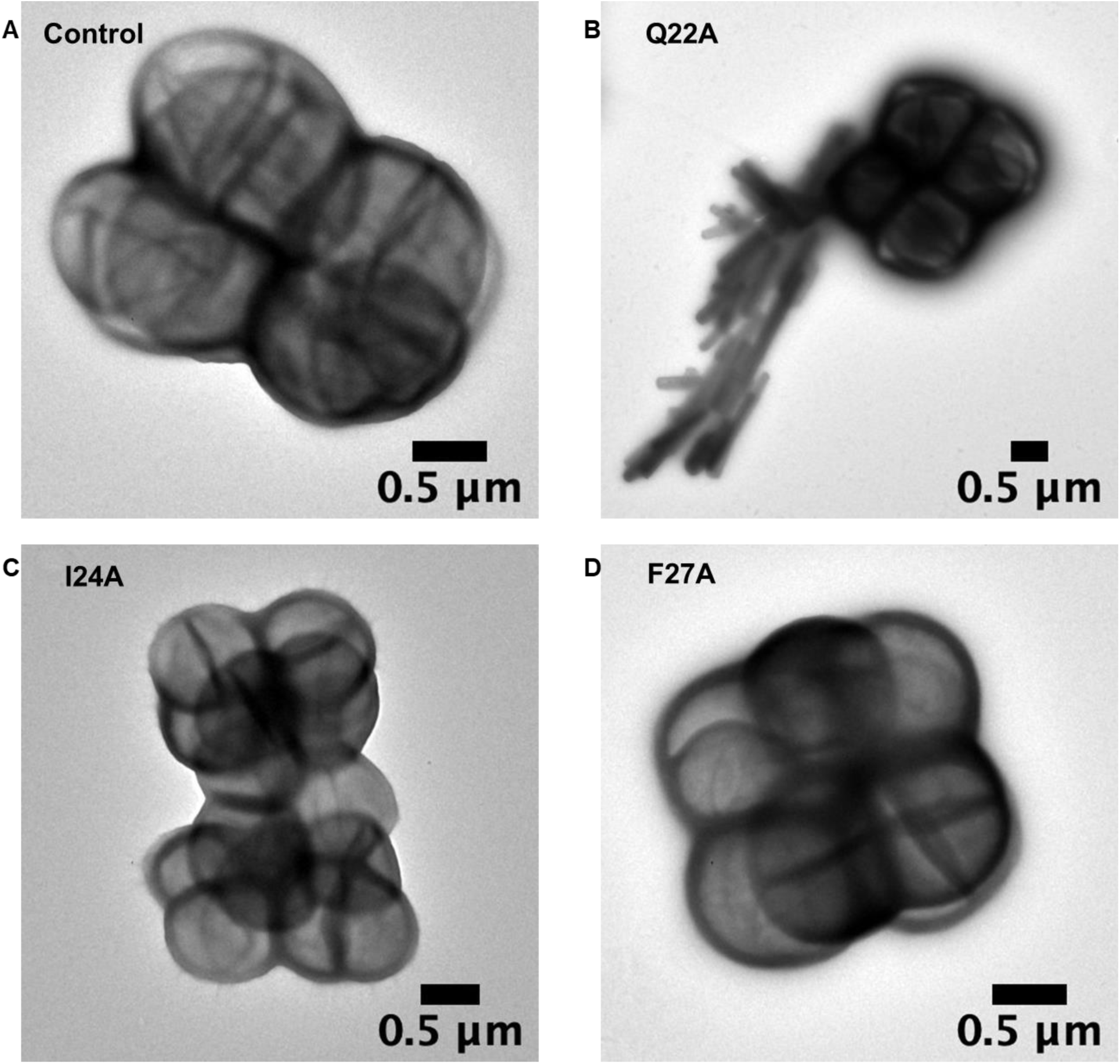
Electron micrographs of *M. luteus* incubated with hLL-37_17-29_ mutants. Transmission electron micrographs of *M. luteus* incubated with hLL-37_17-29_ mutants. (**A**) A control sample containing *M. luteus* with no peptide added. (**B**) The hLL-37_17-29_ Q22A active mutant (50 µM; antibacterial activity is shown in Fig. 1) incubated with the bacteria displayed nano-fiber-like assemblies around and contacting the bacterial cells. The hLL-37_17-29_ I24A (**C**) and F27A (**D**) inactive mutants, incubated at 100 µM with the bacteria, did not assemble.

**Figure S13.**
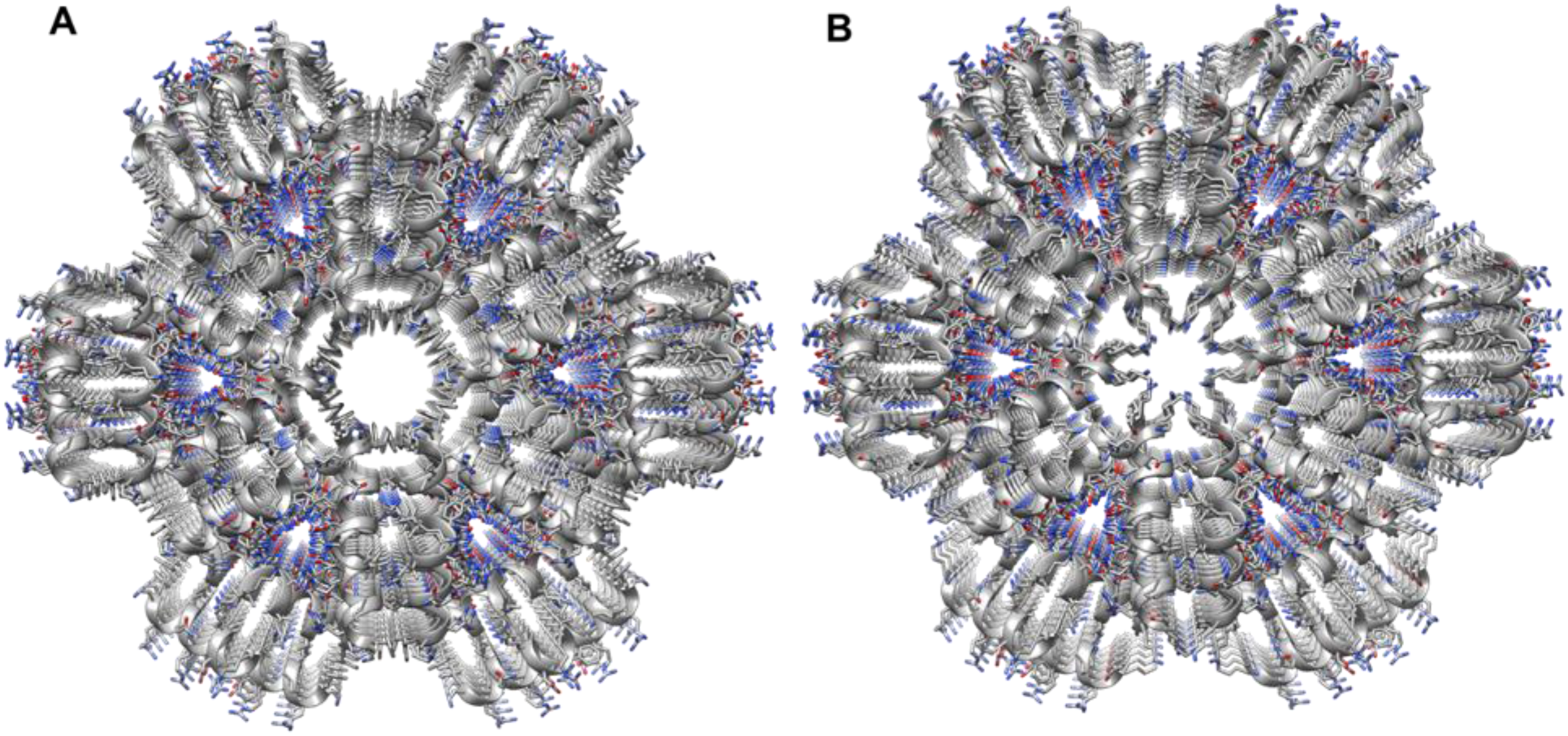
Human and gorilla LL-37_17-29_ share fibrillary architecture but differ in central pore properties. Comparison of human (**A**) and gorilla (**B**) LL-37_17-29_ crystal structures, shown in grey ribbons with side chain shown as sticks colored by atom type (oxygen in red and nitrogen in blue). The view is down the fibril axis, showing the hexametric arrangement and the central pore. The overall structure of the two is highly similar (RMSD of 0.15 Å for the asymmetric unit, comprising two helices and a similar space group and unit cell dimensions (Table S1)). The N-termini of the helices, with Phe17 in hLL-37_17-29_ or Ser17 in gLL-37_17-29,_ and Lys18, lining the central pore. The pore of gLL-37_17-29_ is more occluded, due to the orientation of the lysine residues extending into the pore. In the hLL-37_17-29_ structure, the lysine residues are almost perpendicular to the cross-section of the pore.

**Figure S14.**
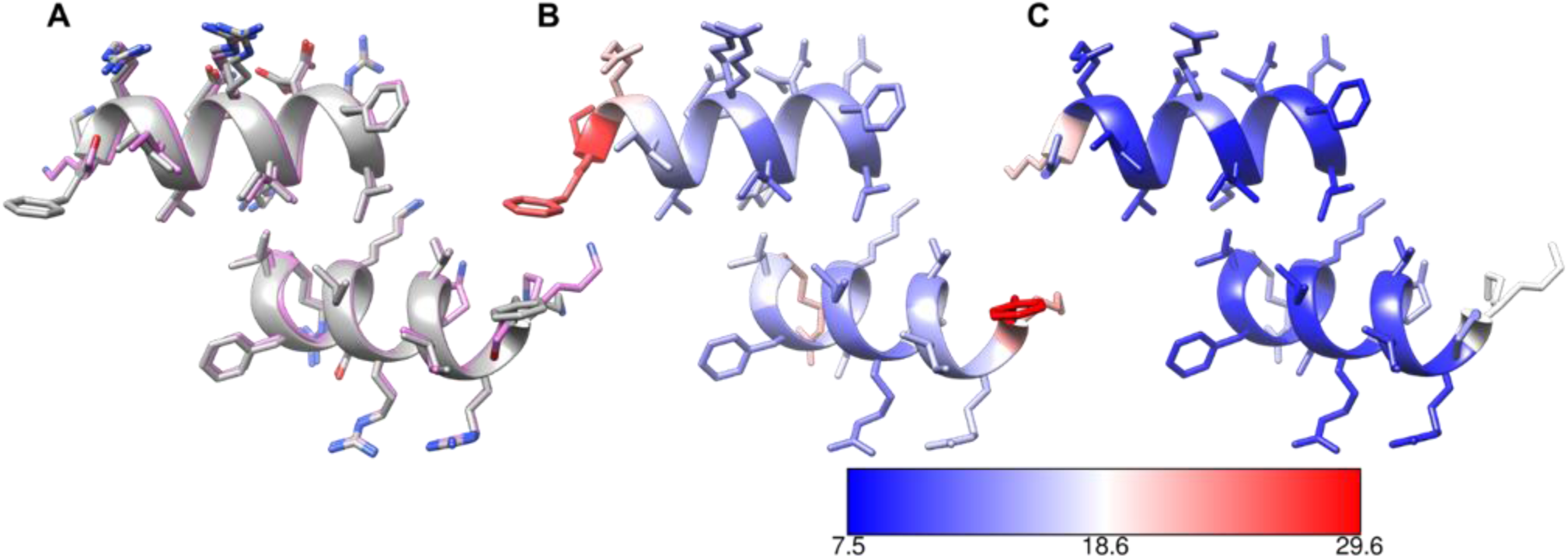
Structural alignment between human and gorilla LL-37_17-29_ and residue flexibility. (**A**) Structural superimposition of the asymmetric unit, comprising two helices, of the human (grey) and gorilla (pink) LL-37_17-29_ crystal structures, revealing a very similar structure with RMSD of 0.15 Å. (**B-C**) Human (**B**) and gorilla (**C**) LL-37_17-29_ in the same orientation as in panel A, colored by average B (temperature) factors per residue, according to the scale bar, with blue-to-red indicating stable-to-flexible (ordered-to-disordered).

**Table S1.**
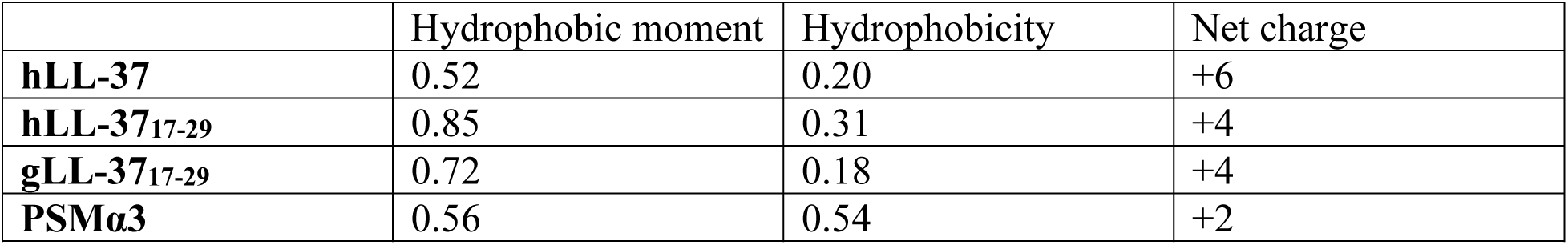
Properties of the amphipathic helices

|  | Hydrophobic moment | Hydrophobicity | Net charge |
| --- | --- | --- | --- |
| **hLL-37** | 0.52 | 0.20 | +6 |
| **hLL-37<sub>17-29</sub>** | 0.85 | 0.31 | +4 |
| **gLL-37<sub>17-29</sub>** | 0.72 | 0.18 | +4 |
| **PSMα3** | 0.56 | 0.54 | +2 |

**Table S2.** Data collection and refinement statistics (molecular replacement)

|  | **Gorilla LL-37<sub>17-29</sub>** | **Human LL-37<sub>17-29</sub>** |
| --- | --- | --- |
| PDB accession code | 6S6N | 6S6M |
| Beamline | ESRF ID23-2 | EMBL P14 PETRA III |
| Date | May 12, 2018 | July 7, 2018 |
| **Data collection** |  |  |
| Space group | P 61 2 2 | P 61 2 2 |
| Cell dimensions |  |  |
| *a*, *b*, *c* (Å) | 41.15 41.15 57.28 | 41.45 41.45 57.47 |
| $\alpha$, $\beta$, $\gamma$ (°) | 90.0 90.0 120.0 | 90.0 90.0 120.0 |
| Wavelength (Å) | 0.873 | 0.976 |
| Resolution (Å) | 57.3-1.1 (1.13-1.10) | 35.9-1.35 (1.39-1.35) |
| R-factor observed (%) | 5.6 (55.2) | 5.6 (78.8) |
| <sup>b</sup>*R*<sub>meas</sub> (%) | 5.7 (59.8) | 6.0 (83.9) |
| *I* / $\sigma$ | 31.1 (3.1) | 21.5 (3.5) |
| Total reflections | 222293 (5198) | 59815 (4470) |
| Unique reflections | 12186 (808) | 6782 (505) |
| Completeness (%) | 99.4 (92.1) | 98.7 (99.8) |
| Redundancy | 18.2 (6.4) | 8.8 (8.9) |
| <sup>c</sup>CC<sub>1/2</sub> (%) | 99.9 (90.4) | 99.7 (89.7) |
| **Refinement** |  |  |
| Resolution (Å) | 35.6-1.1 (1.13-1.10) | 19.5-1.35 (1.39-1.35) |
| Completeness (%) | 99.4 (92.1) | 98.7 (99.8) |
| <sup>d</sup>No. reflections | 10967 (727) | 6103 (452) |
| <sup>e</sup>*R*<sub>work</sub> (%) | 16.0 (47.8) | 23.2 (39.9) |
| *R*<sub>free</sub> (%) | 17.9 (55.9) | 26.8 (39.2) |
| No. atoms | 307 | 300 |
| Protein | 123 (Chain A)<br>128 (Chain B) | 122 (Chain A)<br>147 (Chain B) |
| Water | 56 | 31 |
| *B*-factors |  |  |
| Protein | 11.4 (Chain A)<br>10.5 (Chain B) | 17.2 (Chain A)<br>16.3 (Chain B) |
| Water | 28.8 | 28.1 |
| R.m.s. deviations |  |  |
| Bond lengths (Å) | 0.014 | 0.011 |
| Bond angles (°) | 1.708 | 1.613 |
| Clash score<sup>63</sup> | 1.83 | 1.71 |
| Molprobity score<sup>63</sup> | 0.94 | 0.92 |
| Molprobity percentile<sup>63</sup> | 99<sup>th</sup> percentile | 99<sup>th</sup> percentile |
| Number of xtals used for scaling | One crystals | One crystal |

Values in parentheses are for highest-resolution shell.
<sup>(b)</sup> R-meas is a redundancy-independent R-factor defined in <sup>64</sup>.
<sup>(c)</sup> CC<sub>1/2</sub> is percentage of correlation between intensities from random half-datasets<sup>65</sup>.
<sup>(d)</sup> Number of reflections corresponds to the working set.
<sup>(e)</sup> Rwork corresponds to working set.

**Table S3.**
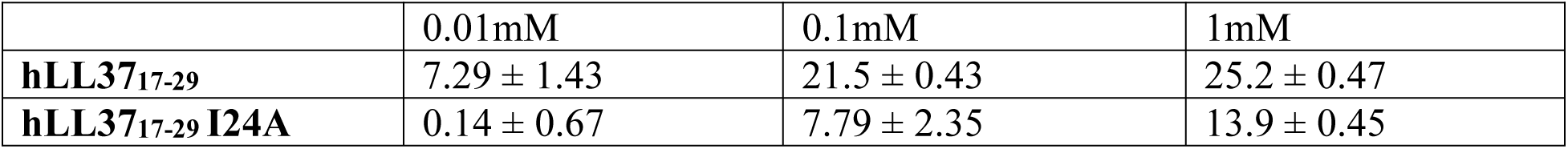
Zeta potential measurements of hLL3717-29 and I24A

**Table S4.**
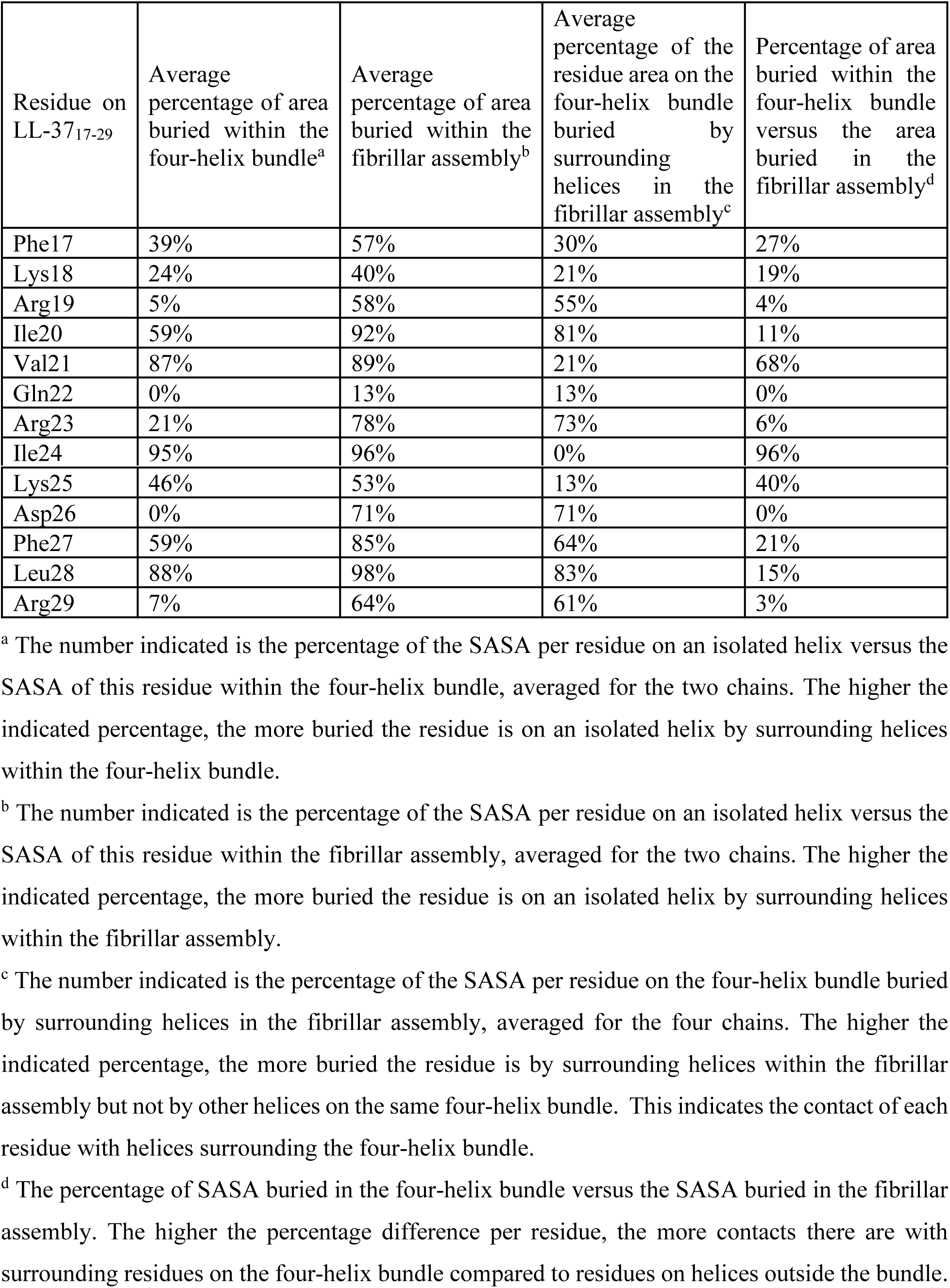
Calculations of the solvent accessible surface area (SASA) per residue in the crystal structure of hLL-37_17-29_

| Residue on LL-37<sub>17-29</sub> | Average percentage of area buried within the four-helix bundle<sup>a</sup> | Average percentage of area buried within the fibrillar assembly<sup>b</sup> | Average percentage of the residue area on the four-helix bundle buried by surrounding helices in the fibrillar assembly<sup>c</sup> | Percentage of area buried within the four-helix bundle versus the area buried in the fibrillar assembly<sup>d</sup> |
| --- | --- | --- | --- | --- |
| Phe17 | 39% | 57% | 30% | 27% |
| Lys18 | 24% | 40% | 21% | 19% |
| Arg19 | 5% | 58% | 55% | 4% |
| Ile20 | 59% | 92% | 81% | 11% |
| Val21 | 87% | 89% | 21% | 68% |
| Gln22 | 0% | 13% | 13% | 0% |
| Arg23 | 21% | 78% | 73% | 6% |
| Ile24 | 95% | 96% | 0% | 96% |
| Lys25 | 46% | 53% | 13% | 40% |
| Asp26 | 0% | 71% | 71% | 0% |
| Phe27 | 59% | 85% | 64% | 21% |
| Leu28 | 88% | 98% | 83% | 15% |
| Arg29 | 7% | 64% | 61% | 3% |

<sup>a</sup> The number indicated is the percentage of the SASA per residue on an isolated helix versus the SASA of this residue within the four-helix bundle, averaged for the two chains. The higher the indicated percentage, the more buried the residue is on an isolated helix by surrounding helices within the four-helix bundle.
<sup>b</sup> The number indicated is the percentage of the SASA per residue on an isolated helix versus the SASA of this residue within the fibrillar assembly, averaged for the two chains. The higher the indicated percentage, the more buried the residue is on an isolated helix by surrounding helices within the fibrillar assembly.
<sup>c</sup> The number indicated is the percentage of the SASA per residue on the four-helix bundle buried by surrounding helices in the fibrillar assembly, averaged for the four chains. The higher the indicated percentage, the more buried the residue is by surrounding helices within the fibrillar assembly but not by other helices on the same four-helix bundle. This indicates the contact of each residue with helices surrounding the four-helix bundle.
<sup>d</sup> The percentage of SASA buried in the four-helix bundle versus the SASA buried in the fibrillar assembly. The higher the percentage difference per residue, the more contacts there are with surrounding residues on the four-helix bundle compared to residues on helices outside the bundle.

## Movie S1. Crystal structure of hLL-37_17-29_ and the assembly of the peptide into fibril

The movie displays the hexameric crystal structure of hLL-37_17-29_ from different orientations and emphases. The helices are shown in grey ribbon presentation, with one representative four-helix bundle colored pink. The movie starts with a view into the fibril axis and zooms in to the representative bundle, showing its side chains. The orientation is then rotated 90° for a view along the fibril axis, and then further rotated along the fibril axis to display the overall helical assembly. The view is again zoomed into the bundle, showing side chains. The orientation is then rotated back to a view into the fibril axis.

## Movie S2. Confocal microscopy imaging of live *M. luteus* incubated with FITC-hLL-37_17-29_

A series of confocal microscopy images was taken for about two hours, at intervals of 10 minutes, and then merged into a movie. The images reveal a rapid accumulation of FITC-hLL-37_17-29_ on the bacteria cells and the formation of peptide foci, indicating rapid aggregation in the presence of *M. luteus*. Depletion of the Hoechst blue signal was observed for bacterial cells localized with the peptide (green), indicating cell rupture and the release of DNA ^66,67^. Bacterial cells with no accumulation of the peptide maintained a strong Hoechst blue signal.

## Materials and Methods

### Peptides and reagents

LL37_17-29_ sequences are derivatives of the human and gorilla cathelicidin antimicrobial peptides CAMP (UniProt IDs P49913 and Q1KLY3, respectively). hLL37_17-29,_ gLL37_17-29,_ their mutants, fluorescein isothiocyanate (FITC)-labeled peptides, and *Staphylococcus aureus* PSMα3 (UniProt ID H9BRQ7) were purchased from GL Biochem (Shanghai) Ltd. as lyophilized peptides, at >98% purity. Thioflavin T (ThT) was purchased from Sigma-Aldrich. Ultra-pure double distilled water (UPddw) were purchased from Biological Industries. Additional reagents and consumable are mentioned below.

Peptide sequences:

LL37: LLGDFFRKSKEKIGKE**<u>FKRIVQRIKDFLR</u>**NLVPRTES

hLL37_17-29_: FKRIVQRIKDFLR

gLL37_17-29_ (F17S): SKRIVQRIKDFLR

FITC-N-hLL37_17-29_: FITC-FKRIVQRIKDFLR

hLL37_17-29_ F17A: **A**KRIVQRIKDFLR

hLL37_17-29_ K18R: F**R**RIVQRIKDFLR

hLL37_17-29_ K18Q: F**Q**RIVQRIKDFLR

hLL37_17-29_ K18H: F**H**RIVQRIKDFLR

hLL37_17-29_ K18A: F**A**RIVQRIKDFLR

hLL37_17-29_ Q22A: FKRIV**A**RIKDFLR

FITC-N-Q22A: FITC-FKRIV**A**RIKDFLR

hLL37_17-29_ I24A: FKRIVQR**A**KDFLR

FITC-N-I24A: FITC-FKRIVQR**A**KDFLR

hLL37_17-29_ I24S: FKRIVQR**S**KDFLR

FITC-N-I24S: FITC-FKRIVQR**S**KDFLR

hLL37_17-29_ I24K: FKRIVQR**K**KDFLR

hLL37_17-29_ I24Q: FKRIVQR**Q**KDFLR

hLL37_17-29_ I24D: FKRIVQR**D**KDFLR

hLL3717-29 F27A: FKRIVQRIKD**A**LR

FITC-N-F27A: FITC-FKRIVQRIKD**A**LR

### Bacterial strains and culture media

*Micrococcus luteus* (*M. luteus*, an environmental isolate) was a kind gift from Prof. Charles Greenblatt from the Hebrew University of Jerusalem, Israel. An inoculum was grown in Luria-Bertani medium (LB), at 30 °C, 220 rpm shaking, 16 h^27^. *Staphylococcus hominis* (subsp. *hominis Kloos and Schleifer S. hominis)* was purchased from ATCC (ATCC® 27844™). An inoculum was grown in brain-heart infusion media (BHI), at 37°C, 220 rpm shaking, 16 h.

### Determination of minimal inhibitory concentrations (MICs)

*M. luteus* and *S. hominis inoculums were* diluted to an OD_600_=0.1. For the MIC experiments, LL37_17-29_ and mutants were dissolved in PBS, and FITC-labeled peptides were dissolved in UPddw. The peptide stock solutions were then diluted into the bacterial media. Control and blank samples contained everything but peptides or everything but bacteria, respectively. Experiments were performed in a sterile 96-well plate and final reaction volume was 100 µl. Bacterial growth (OD_595_) was measured during a 24 h incubation, at 30°C, with 220 rpm shaking, by a plate reader (FLUOstar omega or CLARIOstar, BMG LABTECH). Appropriate blanks were subtracted, and the area under the curve (AUC) was calculated^68^ from the resulting growth curves. MIC values were defined as the minimal concentration of the peptide which yielded less than 20% of the AUC of the control (bacteria with no added peptides). All experiments were performed in triplicates and were averaged. The entire experiment was repeated at least three times on different days, and the mean was calculated from the averaged triplicates of all biological repeats. Error bars represent standard errors of the mean. Two-tailed unpaired t-tests were performed to compare the mean MIC values of tested mutant peptides or derivatives to that of LL37_17-29_.

### Confocal microscopy imaging of peptides interacting with bacteria

*M. luteus* were grown for 16 h and diluted to an OD_600_=0.1. FITC-labeled peptides were dissolved in UPddw, sonicated for 3 min, and then added to the bacteria suspension to a final concentration of 30-150 µM (as indicated in the relevant figure); final reaction volume was 100 µl. Control samples contained everything but the peptide or everything but the bacterium. All samples were incubated, in the dark, at 30°C, with shaking at 220 rpm, for 4 h. Thereafter, 1 ml paraformaldehyde 4% (w/v in PBS) was placed over the samples, for 15 min, at room temperature, in the dark. After fixation, samples were washed three times with fresh PBS, and then treated with Hoechst 33342 (10 mg/ml). All samples were applied to µ-Slides VI 0.4 slides (Ibidi, 80666). Confocal images were acquired using an inverted <u>confocal laser-scanning microscope</u> LSM 710 (Zeiss) equipped with a C-Apochromat 40x water <u>immersion</u> objective lens (NA 1.2) and a Definite Focus unit in an environmental chamber set at 37°C. The laser wavelengths for excitation were 405 nm (Hoechst) and 488 nm (FITC). Brightfield images were collected from the 405 nm laser. Emission was collected sequentially at 410-497 nm for Hoechst and at 493-797 nm for FITC. The pinhole was set for 1 µm. Image processing was done with the Fiji software.

*Time-lapse imaging: M. luteus* bacteria from inoculum were diluted to OD_600_ = 0.1 in LB. Hoechst 33342 (10 mg/ml) was added to bacteria suspension and incubated in the dark, for 10 min, at room temperature. Just before imaging, FITC-LL37_17-29_ was added to the suspensions, to a final concentration of 30 µM. The samples were applied to µ-Slides VI 0.4 slides. Images with a digital resolution of 2048×2048 pixels and 16-bit depth were acquired every 10 min, at room temperature, using a Zeiss LSM 710 confocal microscope. Images were then used to construct a movie using Fiji software. A Gaussian filter (σ=2.0) was applied to the FITC channel for noise removal and brightness and contrast of the images were adjusted^69–71^.

### Thioflavin T (ThT) fluorescence kinetic assay

ThT powder was dissolved in UPddw to a stock solution of 2 mM, vortexed and filtered twice through a 0.22 µm syringe-driven filter unit. Lyophilized PSMα3 peptide was pre-treated as previously described^22^: PSMα3 was dissolved to 1 mg/ml in trifluoroacetic acid (TFA) and hexafluoroisopropanol (HFIP, 1:1v/v), sonicated for 3 min in a sonication bath and left to air-dry in a chemical hood. Dried samples were stored at −20°C. Just before the experiment, PSMα3 was dissolved to 1 mM in UPddw, sonicated for 3 min and immediately transferred to ice. Thereafter, it was diluted to 50 µM with reaction buffer (200 µM ThT, 10 mM sodium phosphate buffer, pH=8, and sodium chloride 150 mM), centrifuged at 12,500 rpm, for 10 min, in a pre-chilled centrifuge (4°C). Lyophilized LL37_17-29_ peptide was dissolved in an identical reaction buffer to 1 mM. Blank solutions contained everything but the peptides. The reaction was carried out in a Greiner Bio-One black 96-well flat-bottom plate, immediately covered with a silicone sealing film (ThermalSeal RTS), and incubated in a plate reader (CLARIOstar or FLUOstar omega, BMG LABTECH) at 25°C, with 220 rpm shaking for 20 sec before each measurement. ThT fluorescence (excitation:438±20 nm; emission: 490±20 nm), was collected for at least 72 h (only 24 h are presented). All measurements were performed in triplicates and the entire experiment was repeated on at least three different days. Readings of blank solutions were subtracted. Error bars represent standard errors of the means.

### Transmission electron microscopy (TEM)

Lyophilized LL37_17-29_ was dissolved in UPddw to a concentration of 1-5 mM and incubated, at 37°C, for several days, as indicated in the individual Figures.

*Imaging the peptides in the presence of bacterial cells*: *M. luteus* was grown for 24 h in LB. Approximately 1.5×10^9^ bacteria cells were washed three times with 10 mM potassium phosphate buffer at pH=7.4. Lyophilized peptides (LL37_17-29_ and its mutants) were dissolved in this same buffer and added to the bacterial pellets, which were re-suspended to a final peptide concentration of 30-150 µM (as detailed in the relevant figures). Samples were then incubated, at 30°C, with 220 rpm shaking, for 4 h.

*Fibril thermostability:* Lyophilized hLL37_17-29_ was dissolved to 2 mM in UPddw and incubated at 37 °C for 3 days. After incubation, samples were moved to a heat block, pre-warmed to 60°C or 80°C (as indicated in the relevant figure), for 10 min. TEM samples were fixed on the EM grid directly from the heat block. The sample heated to 60°C was further incubated at 37°C for 24h and then fixed on the grid.

*TEM grid preparation and visualization*: Samples (4-5 µl) were applied directly onto glow-discharged (easiGlow; Pelco, Clovis, CA, USA, 15 mA current; negative charge; 25 s time) 400 mesh copper grids, with a grid hole size of 42 µm, stabilized with Formvar/carbon (Ted Pella, Inc.). Samples of peptide alone were allowed to adhere for 60 sec, and samples with *M. luteus* were allowed to adhere for 45 sec. Samples were than stained with 1% uranyl acetate solution (Electron Microscopy Science, 22400-1) for 60 sec (peptide alone) or 30 sec (peptide with *M.luteus*), before being blotted with Whatman filter paper. Specimens were examined with a FEI Tecnai T12 G2 electron microscope, at an accelerating voltage of 120 kV, or a FEI Tecnai G2 T20 electron microscope, at an accelerating voltage of 200 kV.

### Cryogenic electron microscopy (cryo-EM)

Lyophilized human and gorilla LL37_17-29_ were dissolved in UPddw to 1-5 mM and incubated at 37°C for 3-10 days (as indicated in the relevant figures). Another sample was prepared from 2 mM hLL37_17-29_ dissolved in 2.7 mM sodium dodecyl sulfate (SDS) (diluted in UPddw from a 40% stock). Of note, 2.7 mM SDS is at sub critical micelle concentration (CMC). Within a temperature-controlled chamber at 100% relative humidity, 3 µl of each sample were applied to a perforated carbon film supported by an electron microscope grid, which were pre-discharged, as described above. After 3 sec, the drop was blotted by Whatman filter paper and liquids were vitrified through rapid plunging of the grids into liquid ethane at its freezing point^72^. Specimens were examined under a FEI Talos 200C high-resolution electron microscope, at an accelerating voltage of 200 kV, using a Gatan 626 cryo-holder. To minimize electron beam radiation damage, the low-dose imaging mode was used. Images were collected digitally by a FEI Falcon III direct-imaging camera and the TIA software, with the help of the “phase plates” (FEI), to enhance image contrast^73,74^.

### Zeta potential measurements

Lyophilized LL37_17-29_ and the I24A mutant were dissolved in UPddw and incubated at 0.01mM, 0.1mM and 1mM concentrations at 37°C for 24 h. E<u>lectrophoretic mobility</u> measurements were performed in 25°C using a Malvern’s Zetasizer Ultra device while samples were measured in folded capillary zeta cell zeta cuvettes (Malvern, DTS1070). Zeta potential was deduced using the Smoluchowski approximation^48^. The cells and the electrodes were washed with ddw three times before and after each sample. Device was equilibrated for 120 sec before each sample and then three consecutive measurements were performed. The presented data is the mean of three consecutive measurements and standard deviation indicates the route mean square of the three measurements.

### Crystallization conditions

Lyophilized human and gorilla LL37_17-29_ peptides were dissolved to 10 mM (17 mg/ml) in double distilled water, vortexed and centrifuged at 14,000 rpm, 4°C, for 10 min. hLL37_17-29_ crystals were grown in a reservoir solution containing 0.2 M sodium acetate, 0.1 M Tris, pH 8.5, and 30% (w/v) polyethylene glycol 4000. gLL37_17-29_ crystals were grown in a reservoir solution containing 0.1 M HEPES, pH=7.5, and 1.4 M sodium citrate. The crystals were grown at 20°C, using the hanging-drop vapor diffusion technique. gLL37_17-29_ crystals were soaked in cryo-protectant solution which contained reservoir solution, 5% (v/v) 2-methyl-2,4-pentanediol (MPD) and 20% glycerol. Crystals were flash-frozen in liquid nitrogen before X-ray data collection.

### Structure determination and refinement

X-ray diffraction of gLL37_17-29_ was collected at the ID23-EH2 micro-focus beamline at the European Synchrotron Radiation Facility (ESRF), Grenoble, France. The wavelength of data collection was 0.873Å. X-ray diffraction of hLL37_17-29_ was collected at the EMBL micro-focused beam P14, at the high brilliance 3rd Generation Synchrotron Radiation Source at DESY: PETRA III, Hamburg, Germany. The wavelength of data collection was 0.976Å. Data indexing, integrating and scaling were performed using XDS and XSCALE^75^. Phases were obtained by molecular replacement using Phaser^76^. For the molecular replacement of gLL37_17-29_, a 13-residue polyalanine idealized helix was used as the search model. For the molecular replacement of hLL37_17-_29, the structure of gLL37_17-29_ was used as the search model. Crystallographic refinements were performed with Refmac5^77^. Further model building was performed using Coot^78^ and illustrated with Chimera (including constructing Movie S1)^79^. The structures of human and gorilla LL37_17-29_ were determined at 1.35 Å and 1.1 Å resolution, respectively. In both structures, there were two peptide chains in the asymmetric unit and water molecules. There were no residues detected in the disallowed region at the Ramachandran plot. Crystallographic statistics are presented in Table S2.

### Sequence alignments and proteolytic digestion prediction

Sequence alignment between LL37_17-29_ and PSMα3 was performed using the MAFFT server^80^. Amino acids are colored by their physicochemical properties^62^. Proteolytic digestion sites in the human LL37 were predicted using the Expasy’s PeptideCutter Tool^81^.

### Calculations of structural properties

The electrostatic potential map, hydrophobicity and B factor scales presented in the figures were created using Chimera^79^. The values of the hydrophobicity scale were according to Kyte and Doolittle^82^. The electrostatic potential was calculated using APBS-PDB2PQR^83^. Helix amphipathicity and the hydrophobic moment (Table S1) were calculated with HeliQuest^84^.

### Solvent-accessible surface area calculations

Solvent-accessible surface areas (SASAs) were calculated using AREAIMOL, with a probe radius of 1.4Å^85,86^, via the CCP4 package^77^. The solvent-accessible buried surface area of each chain in the asymmetric unit was calculated as the area difference between the isolated chain and the chain within the fibril assembly, and is presented as the percentage of the total SASA of the chain. The SASA per residue within different isolated helical assemblies are presented in Table S4.

